# The immunogenomic landscape of cattle

**DOI:** 10.1101/2022.05.31.493931

**Authors:** Ting-Ting Li, Tian Xia, Jia-Qi Wu, Hao Hong, Zhao-Lin Sun, Ming Wang, Fang-Rong Ding, Jing Wang, Shuai Jiang, Jin Li, Jie Pan, Guang Yang, Jian-Nan Feng, Yun-Ping Dai, Xue-Min Zhang, Tao Zhou, Tao Li

## Abstract

Here, we report the *de novo* assembly of a cattle genome using ultra-long-read nanopore sequencing in conjunction with other advanced technologies. The assembled genome contains only 145 contigs (N50 ∼ 74.0 Mb). Compare to the current reference cattle genome ARS-UCD1.2, 154 gaps are closed, and 467 scaffolds are further placed in our assembly. Importantly, except two remained gaps in the T-cell receptor α/δ (TRA/TRD) region, the gene loci of other TRs and immunoglobulins (IGs) are seamlessly assembled and exquisitely annotated. With the characterization of 258 IG genes and 626 TR genes that distributed in seven genomic loci, we illustrate the highest immune gene diversity in mammals to our knowledge. Moreover, the gene structures of major histocompatibility complex (MHC) are integrally depicted with properly phased haplotypes. Thus, our work not only reports a cattle genome with the most continuity and completeness, but also provide a comprehensive view of the complex immune- genome.

## INTRODUCTION

The immune system possesses the biggest source of genetic variation, and the prodigious diversity and complexity of the immune system ensure the host to precisely distinguish non-self from self and to effectively response to the persistent but unpredictable environmental challenges (Schultze and Aschenbrenner, 2021; Sette and Crotty, 2021). At the DNA level, T cells and B cells represent the typical examples of the genetic variations (Imkeller and Wardemann, 2018; Kumar et al., 2018). The somatic rearrangement of the V, (D) and J gene segments from the TR or B cell receptor (BCR, also known as IG) gene loci give rise to millions of different T- and B-cell receptors (Nielsen and Boyd, 2018). Each T- or B-cell, as characterized by the uniquely expressed TR or IG gene, can response to a specific antigen. The TR and IG genes, in together, encode a major part of the immune repertoire (Arunkumar and Zielinski, 2021; Chi et al., 2020). Another example is the MHC gene locus, which contains many genes that involve in the immune defenses and shows the highest diversity among population (Petersdorf and O’HUigin, 2019). Because of the structural complexity of these immunogenetic loci, a comprehensive description of these alleles remains a challenge. The complete assembly and annotation of the immunogenomic loci will provide fundamental and accurate descriptive data for immunology studies. Excitingly, using nanopore sequencing technology, human MHC gene locus was completely assembled and phased with ultra-long reads (Jain et al., 2018a).

The average cost of *de novo* assembly of a genome has dramatically decreased because of the improved next generation sequencing (NGS) technologies such as Illumina(Hu et al., 2021). More importantly, the third-generation sequencing technologies, which can produce long reads that exceeds dozens of kilobases, have led to a paradigm shift in whole-genome assembly, not only in experimental methods but also in algorithms (van Dijk et al., 2018). Pacific Biosciences (PacBio) ‘single-molecule real time (SMRT)’ methods can generate ∼10 Kb long HiFi reads with 99% accuracy (Ardui et al., 2018). Oxford Nanopore Technologies (ONT) recently developed an ultra-long read method that can produce reads with an average length of ∼50 Kb, and the longest reads can be hundreds of kilobases or even over mega-bases (Jain *et al*., 2018a; Nurk et al., 2022). The incredible technical progress has promoted a prosperity of genome assemblies from animals to plants. For human genome, assembly of a centromere on the Y chromosome(Jain et al., 2018b), telomere-to-telomere assembly of specific chromosome (Logsdon et al., 2021; Miga et al., 2020), and a real gapless assembly of all 22 autosomes plus X chromosome (Nurk *et al*., 2022) have been reported recently. These advances provided detailed data and the panoramic view of all genomic variants, especially the immunogenetic diversity of humans.

As one of the most important livestock, cows made important historically contributions and are continuing contribute to the basic and applied immunology (Guzman and Montoya, 2018; Vlasova and Saif, 2021). The basis of vaccination began from the protection against smallpox by inoculation with cowpox (Pead, 2003), and CD205 was firstly identified as dendritic cell marker in cattle (Naessens and Howard, 1993). Cattle also showed privileges in studying human infectious diseases over mouse models such as tuberculosis (Waters et al., 2011) and respiratory syncytial virus (RSV) (Taylor, 2017), as human and bovine share higher similarities in immunity. Moreover, recent investigations of cattle immune responses provided novel insights into the relationships between microbes and host (Gomez et al., 2019). For example, the long third heavy chain complementary determining regions (CDRH3) in cattle has been shown to be capable of rapid elicitation of broad-neutralizing antibodies against Human Immunodeficiency Virus (HIV) (Sok et al., 2017). However, the insufficient understanding of cattle immune system hinders the studies of this important model farm animal. A high-quality reference genome is crucial to facilitate research on cattle immunity.

Besides the current official cattle genome, ARS-UCD1.2, de novo assembly of the cattle genome has been tried with PacBio SMRT method (Rosen et al., 2020). Two genome assembly studies of water buffalo and Simmental cattle have also been reported recently (Heaton et al., 2021; Low et al., 2019). All the above assemblies showed limited genome continuity and completeness, and none of them used ONT ultra-long read method. In this study, we report a new assembly for the cattle genome with a combination of several advanced sequencing methods, in particular, the ONT ultra-long read sequencing technology. Our data significantly surpassed the continuity and accuracy of ARS-UCD1.2, and enabled the gapless assembly and refined annotation of the immunogenetic loci, including TR, IG and MHC.

## RESULTS

### *De novo* Assembly of a Cattle Genome

To assemble a most accurate genome version of cattle, we carried out a whole genome sequencing of female cattle embryonic fibroblasts using ONT ultra-long read sequence technology in conjunction with PacBio circular consensus sequencing (HiFi), Illumina NGS, Hi-C and BioNano optical maps. 237.8 Gb ultra-long reads were produced with an N50 length of 70.4 Kb, and the longest read was 872.5 Kb. The ONT data exhibited a great advantage in read-length compared to that of PacBio and Illumina methods (Figures S1A-C).

Using NextDenovo, we performed the *de novo* assembly of the ultra-long reads (Figure S1D). The assembly comprised only 145 primary contigs with an N50 contig size of 74.01 Mb and a total length of 2.68 Gb. The contigs were then polished and corrected with PacBio HiFi reads and Illumina reads, anchored with BioNano optical maps and Hi-C interaction matrix into a final version genome with excellent continuity (Figure S1E). The N50 scaffold size was improved to 74.72 Mb and the final genome size was 2.71 Gb, and we named our assembly as NCBA1.0 (Figure 1A). To evaluate the completeness and accuracy of the NCBA1.0, we aligned all the reads back to this new assembled genome, and 99% of the whole genome had a minimum coverage of 36 × by ultra-long reads, 4× by PacBio HiFi reads and 64× by Illumina reads (Figures 1B and S1F). The ultra-long reads showed significant low-bias on GC contents of genomic regions compared to that of Illumina reads and HiFi reads, especially at GC-poor regions (Figure S1G). This result exhibited the outstanding performance of ultra-long reads in genome assembly, as heterochromatins consists of AT-rich repeats, especially at the telomeres and acrocentric regions.

**Figure 1.**
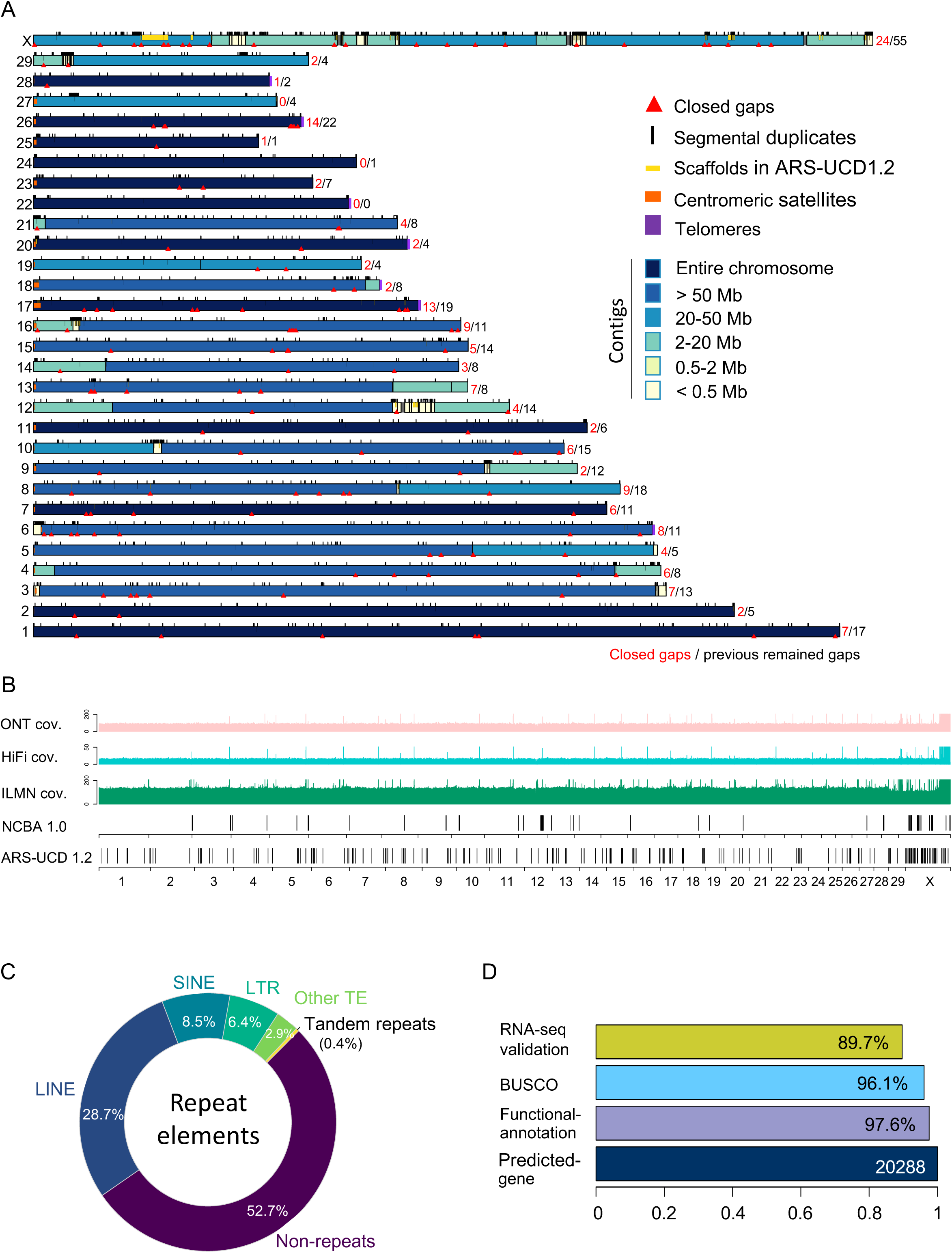
A Global Picture of the *de novo* Cattle Genome Assembly. (A) Ideograph of cattle genome assembly NCBA1.0. Chromosomes composed of a single contig are in dark blue, satDNAs are in orange and telomeres are in purple. Closed gaps and properly placed scaffolds of ARS-UCD1.2 are depicted as red triangles and yellow strips. (B) Read coverage of cattle genome NCBA1.0. Lane 1-3: read coverages by ONT ultra-long reads, PacBio HiFi reads and Illumine NGS reads. Lane 4-5: remaining gaps in NCBA1.0 and ARS-UCD1.2. (C) Composition ratio of repeat elements in NCBA1.0. (D) Gene annotations of NCBA1.0. There were 20288 genes predicted in total, and the ratio of genes with functional annotation, overlapped with BUSCO and validated with RNA-Seq were indicated.

### Gap Filling on the Reference Genome ARS-UCD1.2

The NCBA1.0 assembly showed tremendous sequence integrity compared to the current cattle reference genome, ARS-UCD1.2, and other cattle genomes (Figure S1H). Of the 30 chromosomes, 12 chromosomes were packaged by one single contig (Figure 1A). The q-arms of seven chromosomes were ended with a minimum of 15 Kb (TTAGGG)_n_ telomere repeats, among which five chromosomes (chromosome 17, 20, 22, 26 and 28) were gapless centromere-to-telomere assemblies (Figure 1A). The gap- remaining regions were mainly localized at the acrocentric regions of p-arms and the chromosome X. Further, we assessed whether the remained gaps in ARS-UCD1.2 can be filled using our assembly. The reference cattle genome ARS-UCD1.2 contains 30 chromosomes and 2180 unplaced scaffolds. There are 386 gaps that denoted as Ns and 315 gaps of which were localized on chromosomes. With our assembly, 154 gaps on chromosomes were filled (Figures 1A-B). Additionally, 420 scaffolds and 47 partial scaffolds, with a total length of 24.89 Mb, can be properly placed back in the new genome (Figure S2). The scaffolds placed ranged from Kb to Mb and the largest scaffold was 4.3 Mb in chromosome X.

We further annotated the newly assembled genome. Transposable element repeats (TE) were account for 46.53% of NCBA1.0 and the total ratio of repeat sequences were 47.30% (Figure 1C). 20,288 genes were predicted with an average length of 40.4 Kb, which showed close consistency with bovine and other proximal species (Figures 1D and S3A-F). Functional annotations of the genes among databases, including Benchmarking Universal Single-Copy Orthologs (BUSCO, 95.25% overall alignment rate), KEGG, GO, KOG, NR and Swiss-Prot, showed both high coverage and intersection ratios (Figures S4A-C). We also validated the expression of the predicted genes by RNA-Seq and confirmed the expression of 89.7% of these genes (Figure 1D). These data demonstrated the reliability of the genome annotation of NCBA1.0.

### Immunoglobin Gene Loci Annotation

The assembly of a cattle genome with the most continuity and completeness allowed us to depict the detailed gene structures of the complex immunogenomic loci. We mainly looked into the IG, TR and MHC gene clusters, which localized on six different chromosomes (Figure S5). IG genes or B cell receptors are composed by three gene loci, immunoglobin heavy chain (IGH), lambda chain (IGL) and kappa chain (IGK). All IG-related gene loci were covered with gapless contigs and were well annotated in NCBA1.0, while in genome ARS-UCD1.2, IGH is missing, and IGL region remains six gaps (Figures 2A-D and S5). Detailed gene structure and functionality annotations of IG/TR loci were performed mainly following the IMGT criteria (Figures S6A and S6B). The IGH was 616.0 Kb in size and was located in the end of the q-arm of chromosome 21 (Figures 2A, 2B and S7A). During the maturation of IG, a process called V(D)J recombination takes place. This process combines randomly selected one segment from each of the preexisting variable (V), diversity (D), and joining (J) gene clusters and give rise to the tremendous diversity of IGs on mature B cells (Nielsen and Boyd, 2018). In the NCBA1.0, the IGH locus consists of 48 IGHV genes (11 functional) belonging to 3 IGHV subgroups, and 17 IGHD genes (all functional), 12 IGHJ genes (3 functional) and 10 IGHC genes (8 functional) (Figure 2B). A previous study assembled IHG locus by sequencing seven BAC clones and generated an IHG gene structure containing three tandem [IGHDP-IGHV3-(IGHDv)_n_] repeats (Ma et al., 2016). In contrast, there are only two tandem repeats [IGHDP-IGHV3-(IGHDv)_n_] in IGH locus of NCBA1.0, and the repeat regions as well as the adjacent gene loci were fully covered with multiple ultra-long reads longer than 100 Kb, demonstrating the accuracy and reliability of sequence assembly in our study (Figures S7B and S8 A-B). In addition, compared to the above work, two extra IGHV genes in the V region were identified (Figure 2B). Thus, our data suggested that the IGH locus were organized as: (IGHV)_46_- (IGHDv)_5_-(IGHJ)_6_-IGHM1-[IGHDP-IGHV3-(IGHDv)_n_]_2_-(IGHJ)_6_-IGHM2-IGHD-IGHG3-IGHG1-IGHG2-IGHE-IGHA (Figures 2A and 2B). 10% of cattle IGs contain ultra-long complementarity determining region (CDR3) that were composed of IGHV1-7 and IGHD8 (Deiss et al., 2019; Haakenson et al., 2018). We identified one unique gene locus of IGHV1-7 and one unique IGHD8 in the global IGH gene locus. We renamed these gene loci as IGHV1-6 and IGHD8-2 according to their position in NCBA1.0 (Figure 2B).

**Figure 2.**
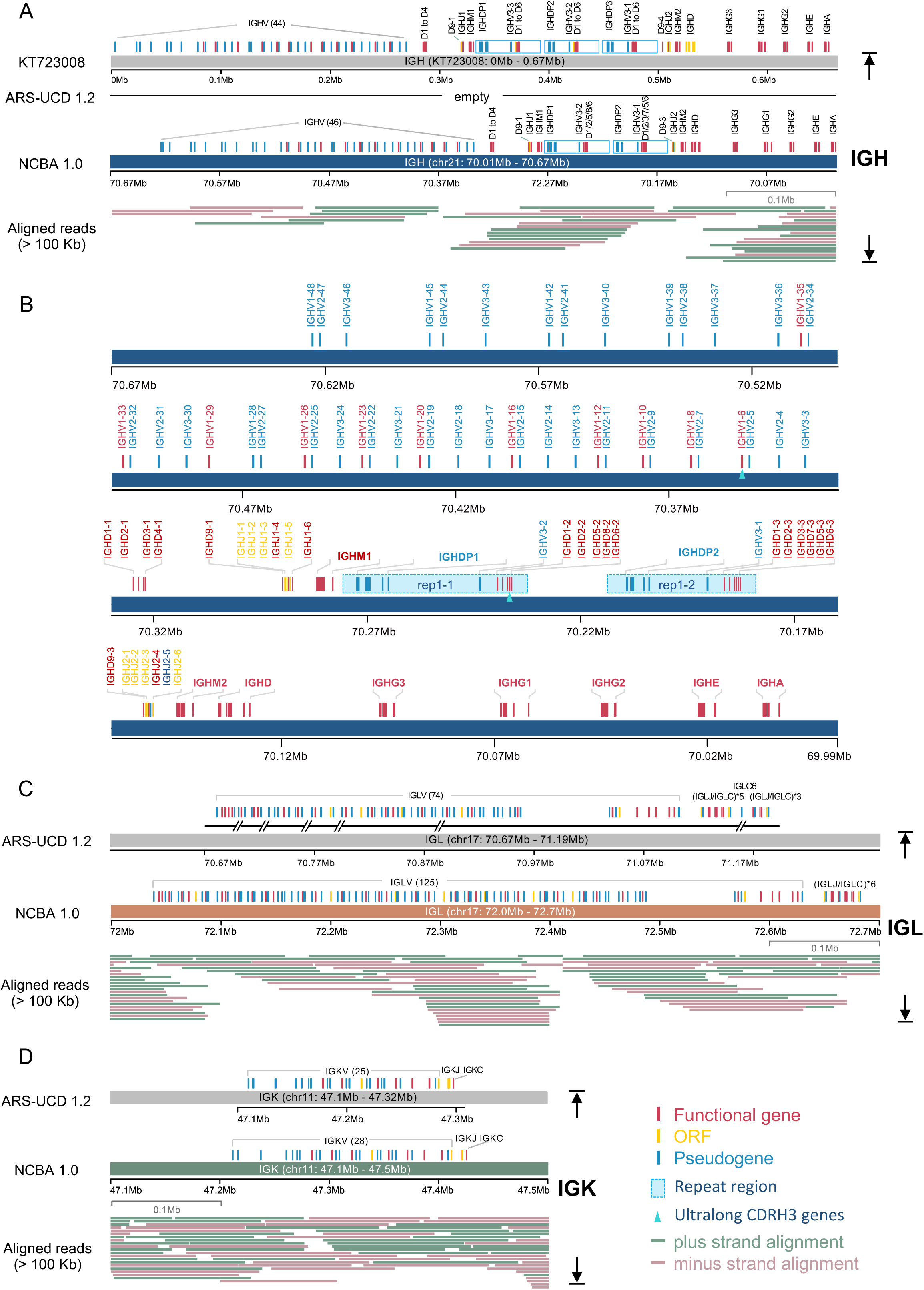
The Cattle Immunoglobin Loci. (A) Genomic organization of IGH locus in KT723008, ARS-UCD1.2 and NCBA1.0. Repeated regions were drawn as blue rectangles, and ONT ultra-long reads that longer than 100 Kb and the mapping to the genomic region were drawn. (B) Detailed diagram of IGH gene structure and annotation in NCBA1.0. Two repeated regions were labeled as blue rectangles (rep1-1 and rep1-2). IGHV1-6 and IGHD8-2 consists of the ultralong CDRH3 were marked with green triangles. (C-D) Genomic organization of IGL and IGK loci in ARS-UCD1.2 and NCBA1.0. Genomic gaps in ARS-UCD1.2 were depicted beneath the genes according to their coordinates and ONT ultra-long reads that longer than 100 Kb and the mapping to the genomic region were drawn.

The IGL locus spanned 643.9 Kb at q-arm of reverse strand on chromosome 17. By filling the remained six gaps in the IGL locus of ARS-UCD1.2, total 125 IGLV genes (37 functional) were annotated, among which 51 IGLV genes were newly recognized (Figures 2C and S9A). We also corrected the repeat numbers of IGLJ-IGLC clusters from nine to six (Figures 2C and S9A-B). The whole IGL genome locus was covered with an average depth of 13 by ultra-long reads longer than 100 Kb, and four ultra-long reads span over the entire IGLC region (Figure S10), giving incontrovertible evidence for the genome assembly and annotation in IGL locus. These results illustrated that the IGL genes are organized as: (IGLV)_125_-(IGLJ-IGLC)_6_.

The IGK is the smallest gene locus of IG and spans 214.3 Kb between 47.2 and 47.4 Mb on chromosome 11 (Figures 2D and S11). IGK consists of 28 V genes (7 functional), 5 J genes (1 functional) and 1 C gene in NCBA1.0 and 3 new V genes were found compared to the previous genome version, ARS-UCD1.2 (Figure S11). Our data suggested that the IGK locus is organized as (IGKV)_28_-(IGKJ)_5_-IGKC.

### T Cell Receptor Gene Loci Annotation

The T cell receptors of cattle are composed by four gene subgroups, TRA, TRB, TRD and TRG, that localized at four genomic loci (Figure S5). TRA and TRD form the most complicated immunogenomic locus together that ranges over 3.3 Mb on the reverse strand of chromosome 10 (Figures 3A and S12). The entire TRD resides within the genetic region of TRA (Figures 3A and 3B). Surprisingly, the V region of TRA/TRD spanned over 3 Mb, which was more than 90% of the total TRA/TRD region. In NCBA1.0, we annotated 281 TRAV genes (148 functional) and 57 TRDV genes (46 functional), while in ARS-UCD1.2, only 183 TRAV genes (85 functional) and 39 TRDV genes (31 functional) were defined. In line with ARS-UCD1.2, the D-J-C cluster consisted of 60 TRAJ genes, 1 TRAC gene, 9 TRDD genes, 4 TRDJ genes and 1 TRDC gene (Figure 3B). Therefore, our data suggested that the TRA(D) genomic structures were organized as [TRA(D)V]_n_-(TRDD)_9_-(TRDJ)_4_-TRDC-TRDV3-(TRAJ)_60_-TRAC.

**Figure 3.**
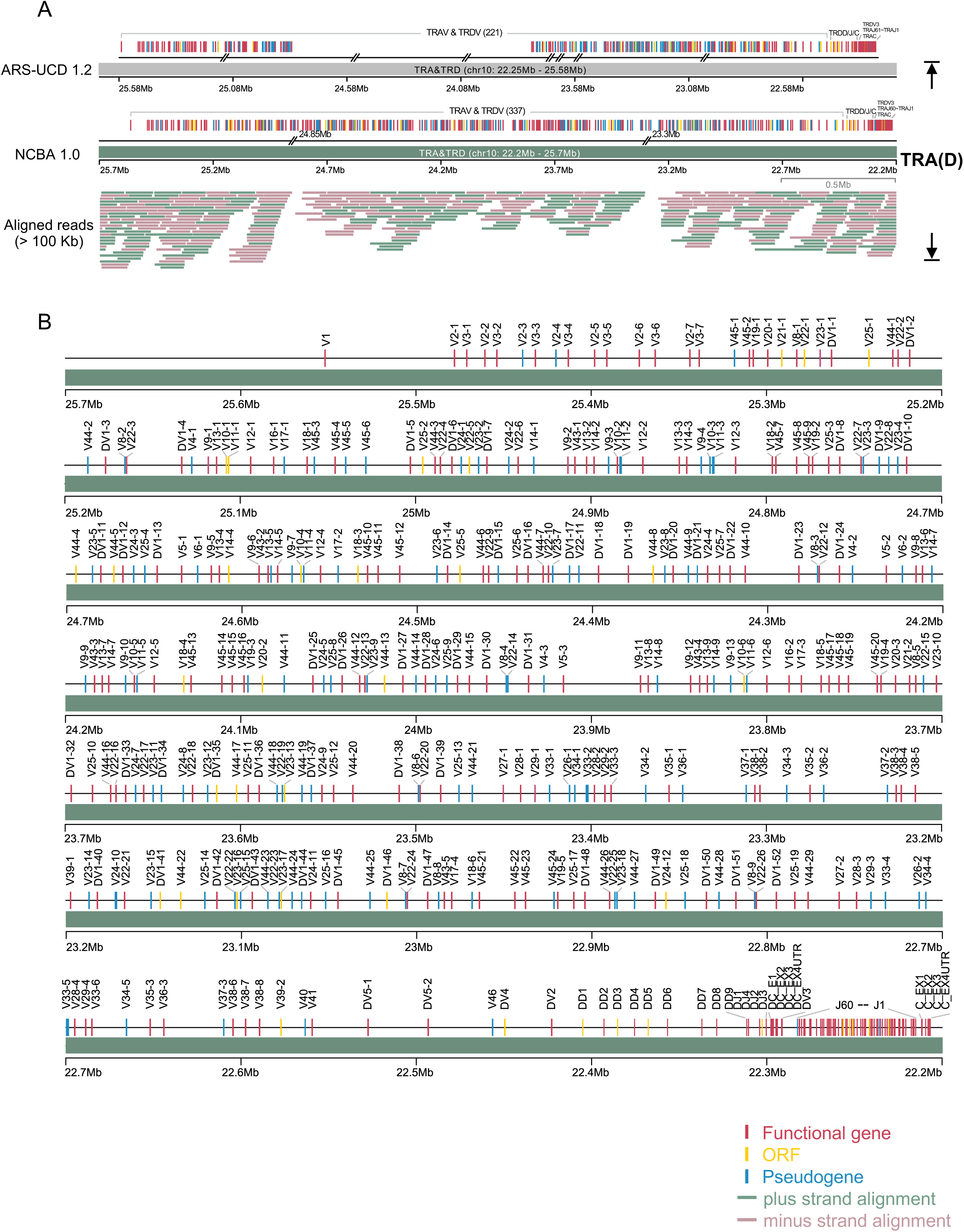
The Cattle TRA/TRD Loci. (A) General organization of TRA/TRD loci in ARS-UCD1.2 and NCBA1.0. Genomic gaps in ARS-UCD1.2 and NCBA1.0 were depicted beneath the genes according to their coordinates. ONT ultra-long reads that longer than 100 Kb and the mapping to the genomic region were drawn. (B) The detailed genetic map of TRA/TRD loci in NCBA1.0. Labels of TRD genes starts with “D” and all TRD genes reside within the TRA genomic region.

The TRB genomic locus was assembled without gap in NCBA1.0. It spanned 667.3 Kb between 105.4 Mb and 106.2 Mb in chromosome 4, and consisted of 153 TRBV genes (87 functional), 3 TRBD genes (all functional), 19 TRBJ genes (15 functional) and 3 functional TRBC genes (Figures 4A and 4B). Our data closed the previously remained two gaps in ARS-UCD1.2 (Figures 4A and S13A). The TRBD, TRBJ and TRBC genes were organized into three tandem D-J-C cassettes, followed by one functional TRBV gene (TRBV30) in inverted orientation (Figure 4B). Thus, the TRB genomic structures were organized as: (TRBV)_152_-[TRBD-(TRBJ)_n_-TRBC]_3_-TRBV30.

**Figure 4.**
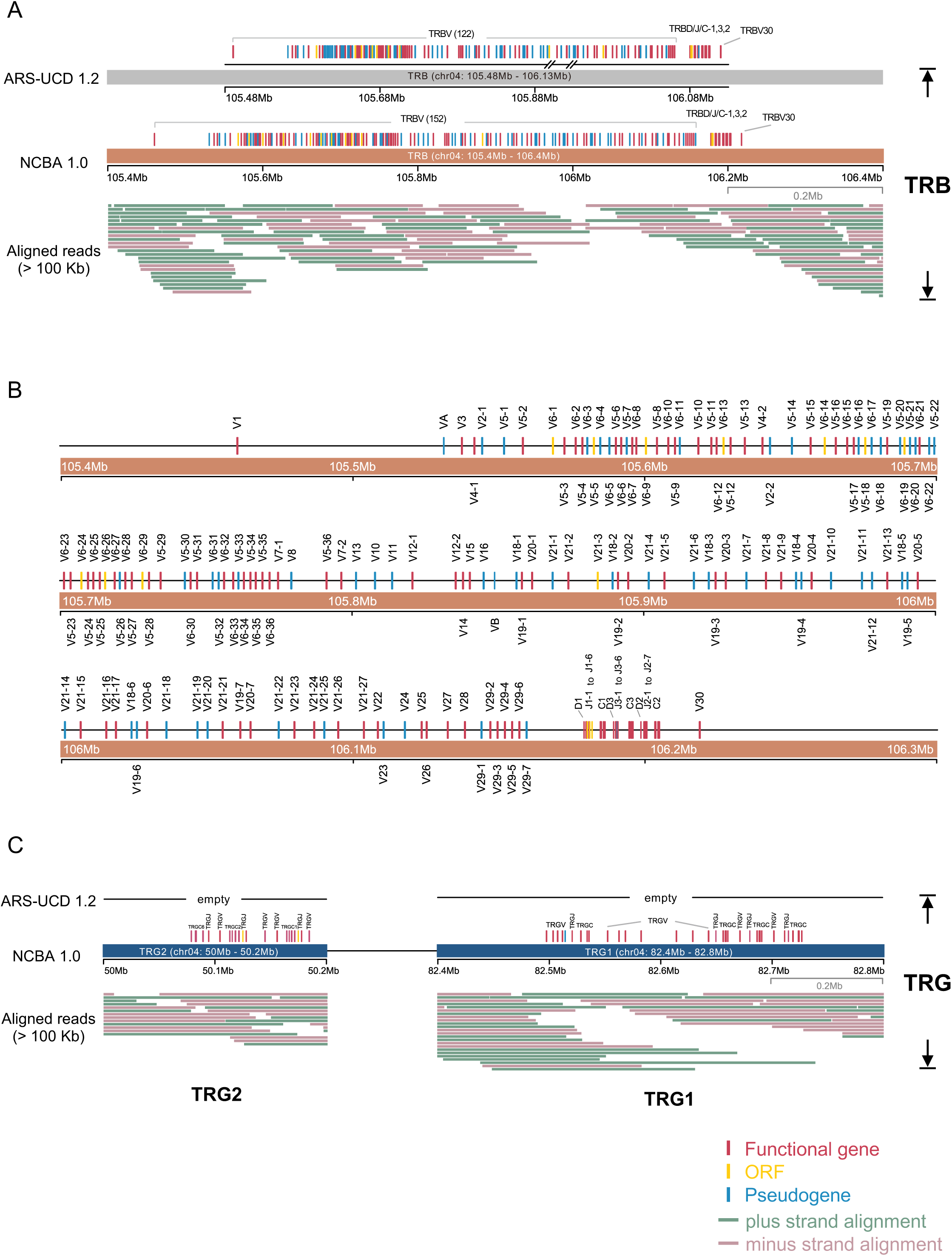
The Cattle TRB and TRG Loci. (A) The genomic organization of TRB locus in ARS-UCD1.2 and NCBA1.0. Genomic gaps in ARS-UCD1.2 were depicted beneath the genes according to their coordinates. ONT ultra-long reads that longer than 100 Kb and the mapping to the genomic region were drawn. (B) The detailed diagram of TRB gene structure in NCBA1.0. (C) The genomic organization of TRG loci in NCBA1.0. ONT ultra-long reads that longer than 100 Kb and the mapping to the genomic region were drawn.

The TRG genes were localized in two separate loci on chromosome 4 which were on different strands and were 30 Mb apart from each other (Figures 4C and S13B). TRG1 spanned 229.3 Kb and comprised four tandem V-J-C cassettes, while TRG2 spanned 106.0 Kb and comprised three tandem V-J-C cassettes (Figure 4C). In total, TRG was consisted of 18 TRGV genes (17 functional), 10 TRGJ genes (8 functional) and 7 TRGC genes (all functional), and TRG genes were organized as [(TRGV)_n_- (TRGJ)_n_-TRGC]_4_ for TRG1 and [(TRGV)_n_-(TRGJ)_n_-TRGC]_3_ for TRG2.

In summary, the cattle genome NCBA1.0 consisted of a total number of 884 IG and TR genes (258 IG and 626 TR) that were localized in seven major loci and distributed in 710 V, 29 D, 116 J and 29 C genes (Table 1). The elaborate annotations of NCAB1.0 greatly enriched the gene sequence diversities compared to that in IMGT database, especially in TR genes (Figure S14). Although there are still two gaps remained in the TRA/TRD region in our assembly, our work provided important data revealing the dramatic diversity of cattle immune repertoire.

**Table 1.** Gene Numbers of Each Immune Locus in NCBA 1.0 Assembly. Functional annotation for each gene of IGs and TRs was performed based on the IMGT criteria (Figure S6), and gene numbers were collected and listed in the table.

| Types | IGH | IGK | IGL | TRA | TRB | TRD | TRG1 | TRG2 | Sum |
| --- | --- | --- | --- | --- | --- | --- | --- | --- | --- |
| V | 48 | 28 | 125 | 281 | 153 | 57 | 14 | 4 | 710 |
| (F/P/ORF) | (11/37/0) | (7/19/2) | (37/80/8) | (148/108/25) | (87/55/11) | (46/7/4) | (13/1/0) | (4/0/0) | (353/307/50) |
| D | 17 | NA | NA | NA | 3 | 9 | NA | NA | 29 |
| (F/P/ORF) | (17/0/0) |  |  |  | (3/0/0) | (6/0/3) |  |  | (26/0/3) |
| J | 12 | 5 | 6 | 60 | 19 | 4 | 5 | 5 | 116 |
| (F/P/ORF) | (3/1/8) | (1/0/4) | (4/0/2) | (53/2/5) | (15/1/3) | (3/0/1) | (5/0/0) | (3/0/2) | (87/4/25) |
| C | 10 | 1 | 6 | 1 | 3 | 1 | 4 | 3 | 29 |
| (F/P/ORF) | (8/2/0) | (1/0/0) | (3/3/0) | (1/0/0) | (3/0/0) | (1/0/0) | (4/0/0) | (3/0/0) | (24/5/0) |
| Sum | 87 | 34 | 137 | 342 | 178 | 71 | 23 | 12 |  |
| (F/P/ORF) | (39/40/8) | (9/19/6) | (44/83/10) | (202/110/30) | (108/56/14) | (56/7/8) | (22/1/0) | (10/0/2) |  |

Table 1. Gene Numbers of Each Immune Locus in NCBA 1.0 Assembly
| Types | IGH | IGK | IGL | TRA | TRB | TRD | TRG1 | TRG2 | Sum |
| --- | --- | --- | --- | --- | --- | --- | --- | --- | --- |
| V<br>(F/P/ORF) | 48<br>(11/37/0) | 28<br>(7/19/2) | 125<br>(37/80/8) | 281<br>(148/108/25) | 153<br>(87/55/11) | 57<br>(46/7/4) | 14<br>(13/1/0) | 4<br>(4/0/0) | 710<br>(353/307/50) |
| D<br>(F/P/ORF) | 17<br>(17/0/0) | NA | NA | NA | 3<br>(3/0/0) | 9<br>(6/0/3) | NA | NA | 29<br>(26/0/3) |
| J<br>(F/P/ORF) | 12<br>(3/1/8) | 5<br>(1/0/4) | 6<br>(4/0/2) | 60<br>(53/2/5) | 19<br>(15/1/3) | 4<br>(3/0/1) | 5<br>(5/0/0) | 5<br>(3/0/2) | 116<br>(87/4/25) |
| C<br>(F/P/ORF) | 10<br>(8/2/0) | 1<br>(1/0/0) | 6<br>(3/3/0) | 1<br>(1/0/0) | 3<br>(3/0/0) | 1<br>(1/0/0) | 4<br>(4/0/0) | 3<br>(3/0/0) | 29<br>(24/5/0) |
| Sum<br>(F/P/ORF) | 87<br>(39/40/8) | 34<br>(9/19/6) | 137<br>(44/83/10) | 342<br>(202/110/30) | 178<br>(108/56/14) | 71<br>(56/7/8) | 23<br>(22/1/0) | 12<br>(10/0/2) |  |

### Phylogenetic Analysis of V Genes

Next, we performed a phylogenetic analysis of V genes across species. Gene diversities statistics of species with detailed IG/TR annotations showed that NCBA1.0 has the highest gene number among known species, especially TR genes (Figure S15). Phylogenetic trees were constructed with all functional IG and TR V genes from human, mouse and cattle (NCBA1.0). Our data showed that all the V genes were well clustered according to their subgroups, except that TRAV genes were clustered into two separate groups (Figure 5A). This result demonstrated the evolutionary conservation of IG and TR gene subgroups among species. The gene number analysis also showed a significant deviation: both human and mouse had much more IG V genes than TR V genes, while cattle showed an opposite tendency that the TR V genes were three times more than IG V genes (Figure 5B). We also evaluated the sequence similarities among the three species. The cattle are more similar to human than mouse by both IG and TR V genes, while human is more similar to mouse by IG V genes and more similar to cattle by TR V genes (Figure 5C).

**Figure 5.**
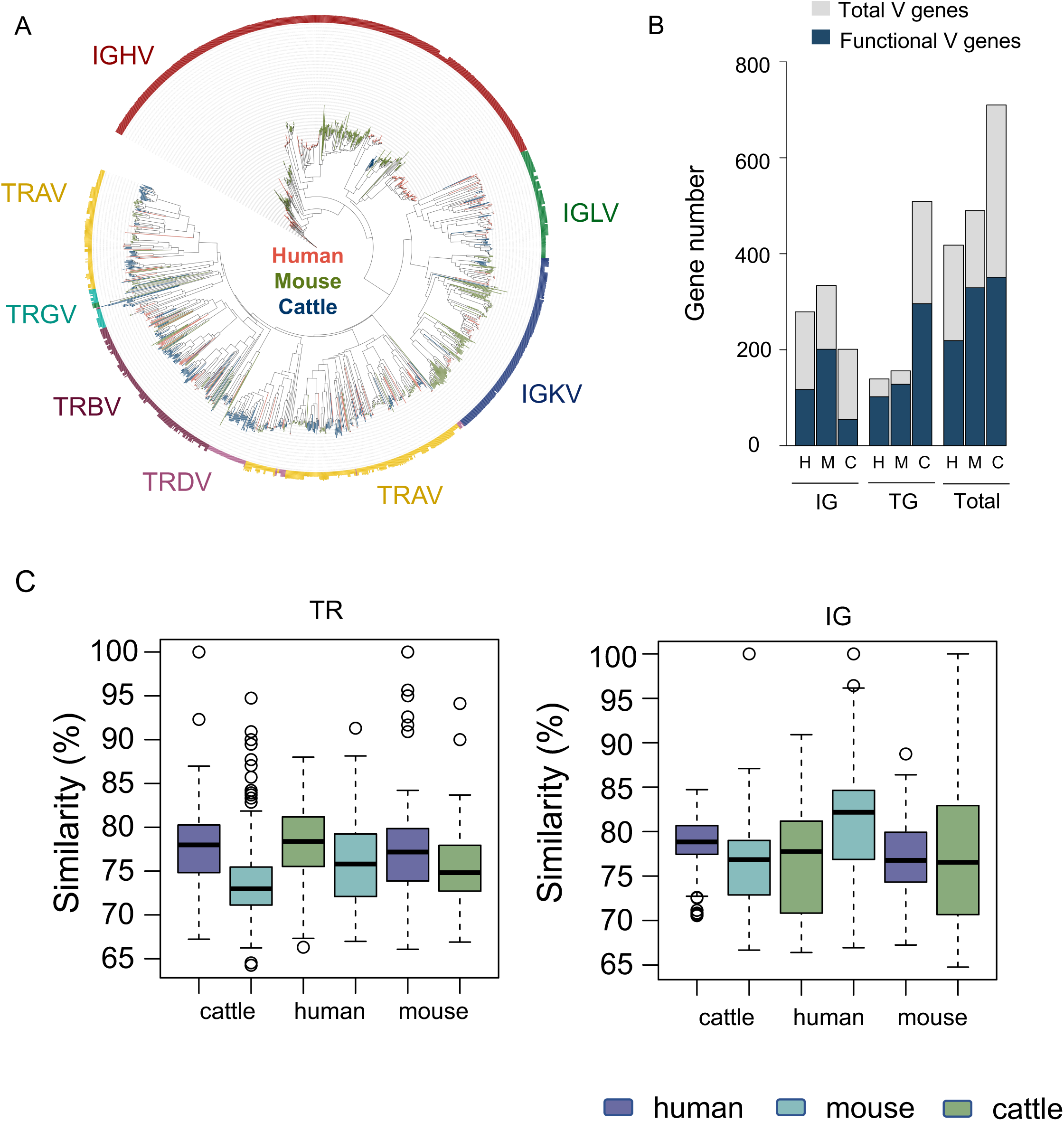
Phylogenetic Analysis of V Genes. (A) Phylogenetic tree of all functional IG and TR V genes. V genes of human, mouse and cattle were merged together and clustered well according to their biological classes. (B) Bar plot of V gene numbers of cattle, human and mouse. (C) Sequence similarities of functional V genes between cattle, human and mouse. Y axis indicates alignment identities of each V gene to the genes of the other two species that share the most similarity.

### MHC Gene Annotation

The MHC plays a crucial role in determining immune responsiveness and is referred as the bovine leukocyte antigen (BoLA) in cattle. In NCBA1.0 assembly, chromosome 23, which contains the BoLA genetic region, was composed by one single contig and the BoLA ranged 3.38 Mb (Figures 6A and S16A). The MHC genes mainly contain two clusters, MHC class I and MHC class II (Petersdorf and O’HUigin, 2019), and by acquiring the seamless sequence of this gene locus, we were able to annotate the MHC genes coordinately. The cattle contain six classical MHC I genes (BoLA 1-6) with high polymorphism and ten non-classical MHC I genes (NC1-10) that show limited polymorphism (Plasil et al., 2022). Classical MHC I genes located at the 3’end of the entire BoLA region. The non-classical MHC I genes NC6-10 were adjacent to classical MHC I genes and genes NC2-5 were 600 Kb upstream away (Figure 6B). For MHC II genes, there were only DQ and DR gene pairs that were organized in adjacent sequential order (Figure 6C), unlike human MHC II that harboring an additional DP gene pair. We also annotated other genes located within the BoLA region, including C2, IL17, CTL4, TNF and TRIM families (Figure 6B and 6C).

**Figure 6.**
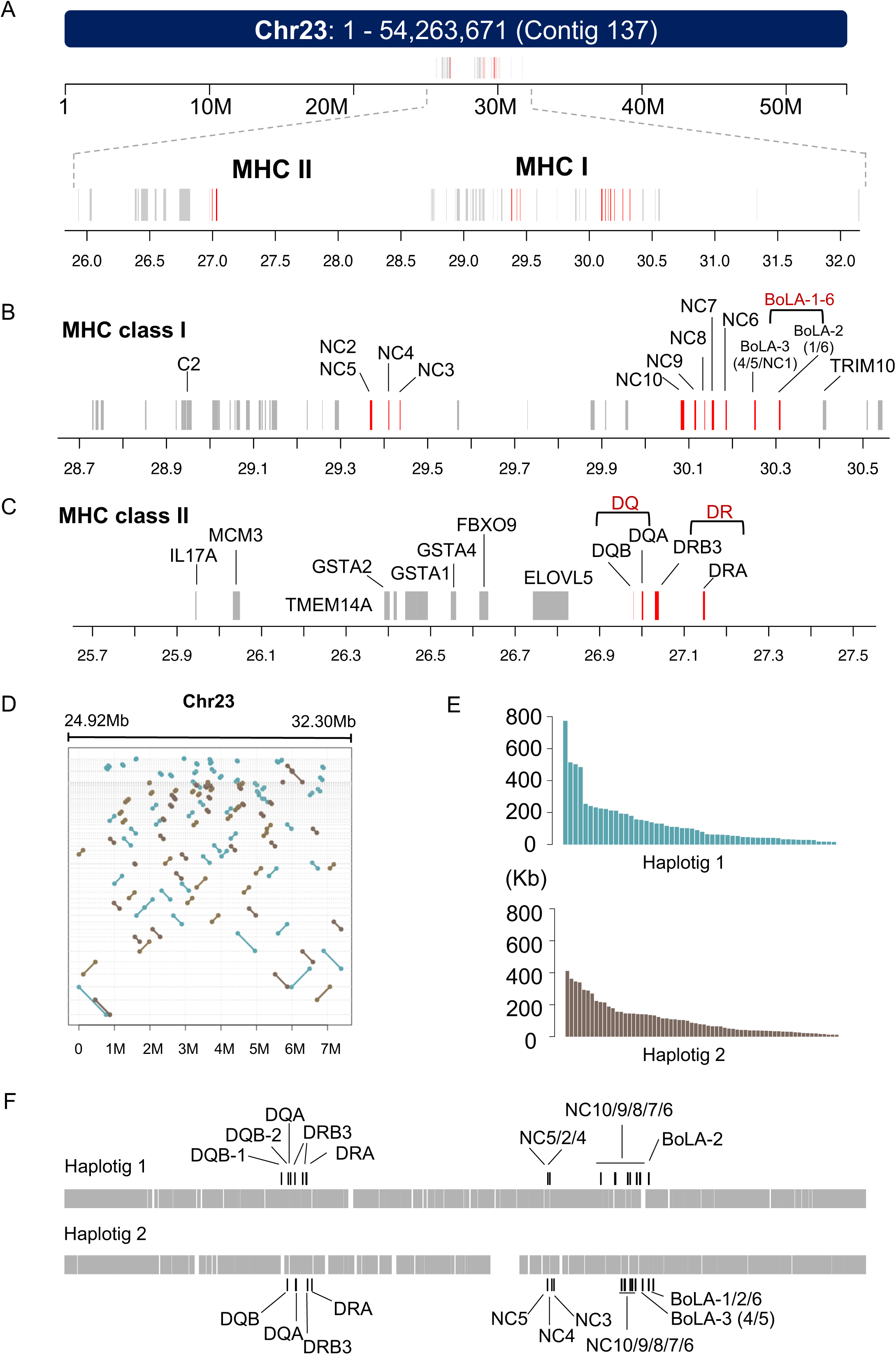
MHC Gene Locus and Haplotyping. (A) Genomic coordinates of MHC locus in chromosome 23. Chromosome 23 consists of only one contig and MHC contains two separate genomic regions. (B-C) Detailed gene organizations of MHC class I (B) and class II (C). MHC genes were labeled in red. (D-F) Haplotyping of MHC genomic region. Genomic locations of two haplotigs within the MHC region (D). Length distributions of two haplotigs (E). Gene organization variation of two haplotigs (F).

To better understand the gene organizations of BoLA region, haplotypes were phased with ONT ultra-long reads combined with heterozygous SNPs called using Illumina data and HiFi reads. The N50 length of haplotype 1 and haplotype 2 were 198.5 Kb and 200.6 Kb, respectively (Figures 6D and 6E). Of the 3.38 Mb BoLA region, 3.17 Mb were successfully assembled into haplotypes and both haplotypes showed high continuity (Figure S16B). The contigs of each haplotype showed delicate differences of gene structures in both MHC I and MHC II (Figure 6F), demonstrating the polymorphism and polygeny of BoLA among individuals. For example, there were two DQB genes within haplotype 1, while only one DQB gene were identified in haplotype 2. Likewise, the classical MHC I genes vary in terms of both gene types and numbers between two haplotypes (Figure 6F). These data highlighted the power of ONT ultra-long technology in resolving haplotypes of huge intricate gene clusters. Taken together, the BoLA was assembled and phased over its full length in a diploid cattle genome for the first time.

### Characterization of Telomere Repeats and Satellite DNA (satDNA)

Heterochromatin, such as centromeres, normally contain long arrays of tandem repeated DNA sequences, known as satDNAs (Escudeiro et al., 2019a; Miga, 2019). Similarly, the telomeres are genomic regions at the end of chromosomes and consist of highly repeated hexanucleotide sequence, TTAGGG (Shay and Wright, 2019). The genomic structure of these repetitive regions in cattle remained elusive due to the lack of investigations of cattle genome with advanced ultra-long sequencing techniques. Thus, we further looked into the telomeres and acrocentric satDNA repeats of the cattle genome NCBA1.0. We identified 1429 ultra-long reads with minimum length of 50 Kb that contains telomere TTAGGG tandem repeats and successfully assembled telomere regions of q-arms of seven chromosomes, and the telomere arrays ranged dozens of kilobases and the longest one spanned over 70 Kb (Figure 7A). As all autosomes of cattle genome are acrocentric (Frohlich et al., 2017), no telomere of p-arms was assembled due to the genomic structures of satDNA arrays in the centromere, which located right beside the telomeres.

**Figure 7.**
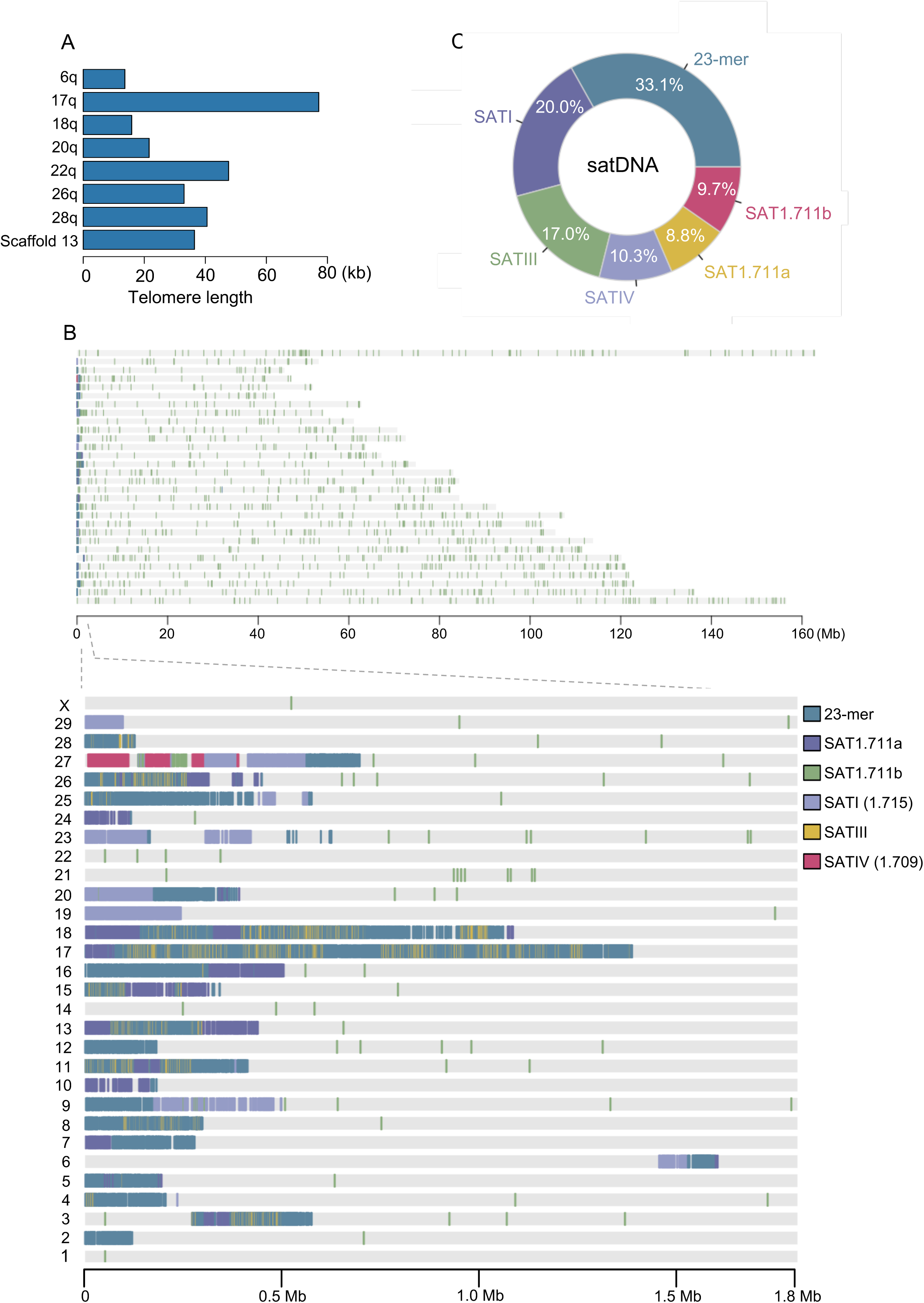
Telomere Length and satDNA Distributions. (A) Lengths of telomere repeats in q-arms of seven chromosomes. (B) Genome-wide distributions of cattle satDNAs. satDNAs mainly localized within the acrocentric regions except sat1.711b. (C) Composition chart of satDNAs in NCBA1.0 assembly.

We next analyzed the distributions of satDNAs in NCBA1.0. Eight types of cattle satDNAs have been reported previously (Ashari et al., 2012) (Escudeiro *et al*., 2019a; Escudeiro et al., 2019b). In our data, FIVE types of satDNAs, SATI, SATIII, SATIV, SAT1.711a and SAT1.711b, constituted the majority of the satDNA regions (Figure 7B). We also identified thousands of copies of a 23-mer arrays, which accounted for the highest abundance of all the satDNA repeats (Figures 7B and 7C). Interestingly, one type of the satDNAs, SAT1.711b arrays, were sparsely distributed along the whole genome, while other satDNAs were located mainly within the acrocentric regions of p-arms (Figures 7B and S17). Collectively, we provided important information for telomeric and satDNA array organizations of cattle genome.

## DISCUSSION

Sequencing-technology progress, especially the emerging of ONT ultra-long sequencing, has achieved great success in the complete human genome assembly, T2T- CHM13, which obtained gapless sequences for all chromosomes except Y (Nurk *et al*., 2022). As the immune system possesses the biggest source of genetic variation, depicting the immune genomic loci was almost impossible several years ago, let alone population genomic diversity studies of the immune system. A completely assembled genome, or at least completely assembled immune genomic loci, is the cornerstone for the in-depth understanding of the immune system of a given species. In this study by taking advantage of the ONT ultra-long sequencing method, we presented a new cattle genome assembly that exhibited remarkable improvement over existing assemblies. We illustrated the advances of our new assembly in gap filling and scaffold assignment. Importantly, we delineated the complex genomic structures of TR, IG and MHC loci, which provides fundamental immunogenomic data for further immune studies in cattle.

Our work affords the most precise roadmap of cattle immune-genome up to now. We corrected the tandem repeat regions within the IGH locus from three to two, and filled 13 gaps within the IG and TR genomic regions. 710 V genes were annotated in NCBA1.0, while only 524 V genes are currently collected in IMGT database (Table 1 and Figure S14). The cattle MHC region was also seamlessly packaged and properly phased, which is the first intact MHC assembly beside human to our knowledge (Jain *et al*., 2018a).

Cattle showed distinct characteristics in both IG and TR immune repertoire. For IGH genes, there were only 48 V genes, which was significantly fewer than human and mouse. The relatively low diversity of IG in cattle was likely compensated by the ultralong CDRH3s that can be found in approximately 10% of the immunoglobins (Haakenson *et al*., 2018). CDRH3s allow cattle antibodies to bind a wider range of antigens and was showed to play a key role in neutralizing HIV spike protein during the immune response (Deiss *et al*., 2019; Sok *et al*., 2017; Wang et al., 2013). These cattle ultralong CDRH3s almost exclusively used the same V gene segment (IGHV1-7) that contains an eight-nucleotide duplication “TACTACTG” at its 3’ end, and the same D gene segment (IGHD8-2) that is known as the longest D gene (Deiss *et al*., 2019). Both IGHV1-7 and IGHD8-2 gene loci were clearly depicted in NCBA1.0. Based on our data, the V-D rearrangements between these two loci can be further investigated, for instance, their histone modifications and chromatin accessibility.

Cattle possess the highest TR gene diversities among all species that were annotated with detailed V(D)J gene structures (Figure S15). We identified 509 TR V genes in NCBA1.0, which was three times more than human TR V genes. The reason why cattle genome contains much more TR V genes than IG V genes remains indistinct. This is probably related with the gut-associated mucosal tissue that contacts with a great diversity of food and microbial antigens, as mucosal T cells play a central role in distinguishing dietary proteins and commensal bacteria from harmful pathogens (Chase and Kaushik, 2019). It is worth mentioning that the remarkably abundant TR repertoire in cattle may serve as a natural resource pool for the screening of specific TRs with extraordinary therapeutic activity against human diseases, such as cancer. Moreover, cattle have a large proportion of γδ T cells that showed regulatory and antigen-presenting functions (Guzman et al., 2012). Further studies of cattle T cells shall shed light on the understanding of γδ T cells, which remains elusive in humans due to their low abundance (Guzman et al., 2014) .

Limited studies in cattle centromeres and telomeres have been done (Escudeiro *et al*., 2019b; Ilska-Warner et al., 2019). Despite the telomere length is associated with productive lifespan and fitness. In this study, we captured telomere repeats in seven chromosomes and for the first time precisely evaluated the telomere length in cattle. Evaluation of satDNAs based on the new assembly revealed sequence ranges and composition ratio of satDNAs, as well as the specific patterns of how different satDNAs were organized that have never been revealed before.

In summary, the new assembly, NCBA1.0, represented a more complete and accurate reference of cattle genome, as well as the immune genome, thereby facilitating further investigations of the immune system in cattle, and perhaps other mammals. These data can be a blueprint for the final gapless telomere-to-telomere cattle genome assembly in the near future.

## ACKNOWLEDGEMENT

This work was supported by grants from the National Key Research and Development Program of China (No. 2020YFA0707702 to Tao Li and No. 2020YFA0707703 to T.Z.) and China National Natural Science Foundation (No. 81925017 to Tao Li, No. 81872153 to T.Z.).

## AUTHOR CONTRIBUTIONS

T.L., T.-T.L. and T.Z. conceptualized and designed the project. T.X., Z.-L.S., M.W., and F.-R.D. prepared the cells. T.-T.L, J.-Q.W. and T.X. analyzed the data. T.-T.L, J.-Q.W., T.X., H.H., S.J., J.L., J.W., G.Y., J.-N.F., Y.-P.D. and J.P. annotated the IG, TR and MHC genes. T.L., T.-T.L. T.X. and X.-M.Z. wrote the manuscript with the help of all authors.

## MATERIALS AND METHODS

### Sample Collection and Genomic DNA Isolation

The holstein cattle of high genetic merit were mated to produce an elite fetus that was recovered at day 60. The cattle fibroblast cells were isolated from this fetus via disaggregation of all tissue excluding the viscera and limbs, and cultured in Dulbecco’s Modified Eagle’s Medium (DMEM; Gibco, Grand Island, New York, USA) supplemented with 10% fetal bovine serum (FBS; Gibco, Grand Island, New York, USA) at 37.5 °C in an atmosphere of 5% C O2 and humidified air. The genomic DNA was extracted using the QIAGEN Genomic-tips kit (QIAGEN, Valencia, CA, USA) according to the manufacturers’ instructions.

### Ultra-long Library Construction and Sequencing

To obtain ultra-long reads, only the large DNA fragments were recovered with BluePippin, followed with end repair and dA-tailing (NEBNext Module, MA, USA). After careful purification, the adapter ligation was performed with SQK-LSK109 ligation kit (Oxford Nanopore Technologies, Oxford, UK) and the final product was quantified by fluorometry (Qubit) to ensure >500 ng DNA was retained, and sequenced on the Oxford Nanopore PromethION platform. ONT ultra-long reads were generated by Grandomics Biosciences company and only reads with a minimum mean quality score of 7 were kept for the following assembly.

### Hi-C Library Construction and Sequencing

The Hi-C experiment was performed exactly following the *in situ* Hi-C method (Rao et al., 2014). Briefly, the cross-linked cells were lysed and digested with MboI, filled with biotin-dATP, ligated with T4 DNA ligase and reverse crosslinked. Then the biotin-labeled DNA was enriched and sequenced with Illumina sequencing platforms following the manufacturer’s instructions. The read mapping, quality control and matrix building were performed with HiC-Pro (Servant et al., 2015).

### De Novo Genome Assembly

The de novo assembly of ONT ultra-long reads was performed with NextDenovo (https://github.com/Nextomics/NextDenovo). The reads were first self-corrected to generate consensus sequences with NextCorrect module and then assembled into preliminary assembly with NextGraph module. To correct the preliminary assembly, the original ONT reads and PacBio CCS reads were mapped with minimap2 (Li, 2018) and corrected with Racon (Vaser et al., 2017) with default parameters for three iterations. Then the Illumina reads were used to polish the corrected assembly with NextPolish (Hu et al., 2020) for 4 iterations to generate the final polished genome assembly. The polished assembly was used as reference for the de novo assembly of BioNano data to generate scaffolds. For Hi-C data, LACHESIS (Burton et al., 2013) was used to cluster, order and direct the scaffolds to generate the final chromosomal level genome assembly.

### Assembly Evaluation

BUSCO 3.1.0 (-l mammalia_odb9 -g genome) (Manni et al., 2021) was used to evaluate the genome completeness based on included gene numbers, and CEGMA v2 (Parra et al., 2007) was used to assess the assembly based on included eukaryotic protein core families with default parameters. Sequence accuracy was assessed by total number of homozygous SNPs identified by Illumina reads mapped to the assembly. Exogenous pollution was assessed based on the distributions of GC-depth and reads coverage.

### Genome Annotation

For repeated sequences, TRF (Benson, 1999) was used to identify tandem repeats and RepeatMasker (Tarailo-Graovac and Chen, 2009) was used to identify transposon-based elements. For gene structures, PASA (Haas et al., 2003)was used to predict gene coordinates based on Illumina RNA-Seq data, GeMoMa (Keilwagen et al., 2018) was used to predict gene coordinates based on protein sequences of proximal species, and GeneMark-ST (Lomsadze et al., 2005) was used to predict genes from de novo. The three gene sets were integrated into an initial gene set with EVM (Haas et al., 2008) and finally to a clean gene set by removing (http://transposonpsi.sourceforge.net/). For further gene function annotation, the protein sequences of the predicted gene set were searched against several databases to predict their functions, including Non-Redundant Protein Database (NR), Kyoto Encyclopedia of Gene and Genome (KEGG), Eukaryotic Orthologous Groups of protein (KOG), InterProScan GO database, and Swiss-Prot.

### Genome Comparison and Gap filling

Genome sequence comparison between ARS-UCD1.2 and the new assembly was performed with LASTZ (Harris, 2007) at chromosomal level. To fill the gaps of ARS- UCD1.2, 10 kb sequences up and downstream of the gap sites in chromosomes were fetched and aligned to the new assembly with minimap2 (Li, 2018). Only alignments with >90% identity were kept and the alignment results of pair of gap sequences were manually checked to ensure the gap loci were within one contig. The unplaced scaffolds were first split up at gap loci and then aligned to the new assembly with minimap2. The scaffolds were reported if > 50% alignment identity and located within one contig.

### Telomeres and satDNA Annotation

To get the loci and lengths of telomeres, all short tandem repeats were identified with TRF (Benson, 1999) with in the new assembly. Then short tandem repeats of TTAGGG were identified and only these located at the end of chromosomes were kept as telomeres. To annotate satDNAs, 57 nucleotide sequences belong to 8 satDNA classes were collected from NCBI and a blast database were built containing these 57 sequences. The sequence similarities among these 57 sequences were analyzed with blast (Johnson et al., 2008). Then all short tandem repeats were identified with TRF, and their pattern sequences were cleaned up to a fasta file and then blasted against the satDNA database. The alignments were filtered with >80% sequence identity and the locations and copy numbers were merged from TRF results.

### IG and TCR Gene Annotation

All cattle IG and TCR gene sequences were downloaded from IMGT database (Lefranc et al., 2015). The gene sequences were aligned to the new assembly with bowtie2 (Langmead and Salzberg, 2012), and the alignment results were merged and manually checked according to their subgroups (IGH, IGK, IGL, TRA, TRB, TRD and TRG) to make sure the gene clusters were seamlessly assembled. For each locus, all candidate variable (V), diversity (D), joining (J), and constant (C) genes were manually annotated according to the following criteria (Figure S6). Manual annotations were validated by four irrelevant people.

### Phylogenetic Analysis of V Genes

Functional IG and TR V gene sequences of human and mouse were downloaded from IMGT database (Lefranc *et al*., 2015). Functional cattle V genes were retrieved from NCBA1.0. Multiple sequence alignment was performed with Clustal Omega (Larkin et al., 2007), and the outputs were visualized using the Interactive Tree of Life software (Letunic and Bork, 2019).

### MHC Gene Annotation and Haplotyping

Totally 713 BoLA gene allele sequences were downloaded from IPD-MHC database (https://www.ebi.ac.uk/ipd/mhc/). The BoLA gene sequences were aligned to the new assembly with bowtie2 (Langmead and Salzberg, 2012), and the genomic location and order of MHC Class I/II genes were manually conformed with alignments > MAPQ 20. For haplotyping, PacBio HiFi reads that mapped to MHC region were retrieved and used for variant calling, followed with SNP genotyping with WhatsHap (Patterson et al., 2015). Then the genotyped reads were separately retrieved and assembled into haplotypes with Canu pacbio-hifi mode (Koren et al., 2017).

### Statistical Analysis

Quantification methods and statistical analysis for each of the separate and integrated analyses are described and referenced in their respective Method Details subsections.

### Data and Code Availability

The relevant data reported in this paper are available from the corresponding authors upon reasonable request. The related codes and figures for reproducible research are stored at GitHub (https://github.com/TintingLi/cattleGenome).

## SUPPLEMENTAL INFORMATION TITLES and LEGENDS

**Figure S1.**
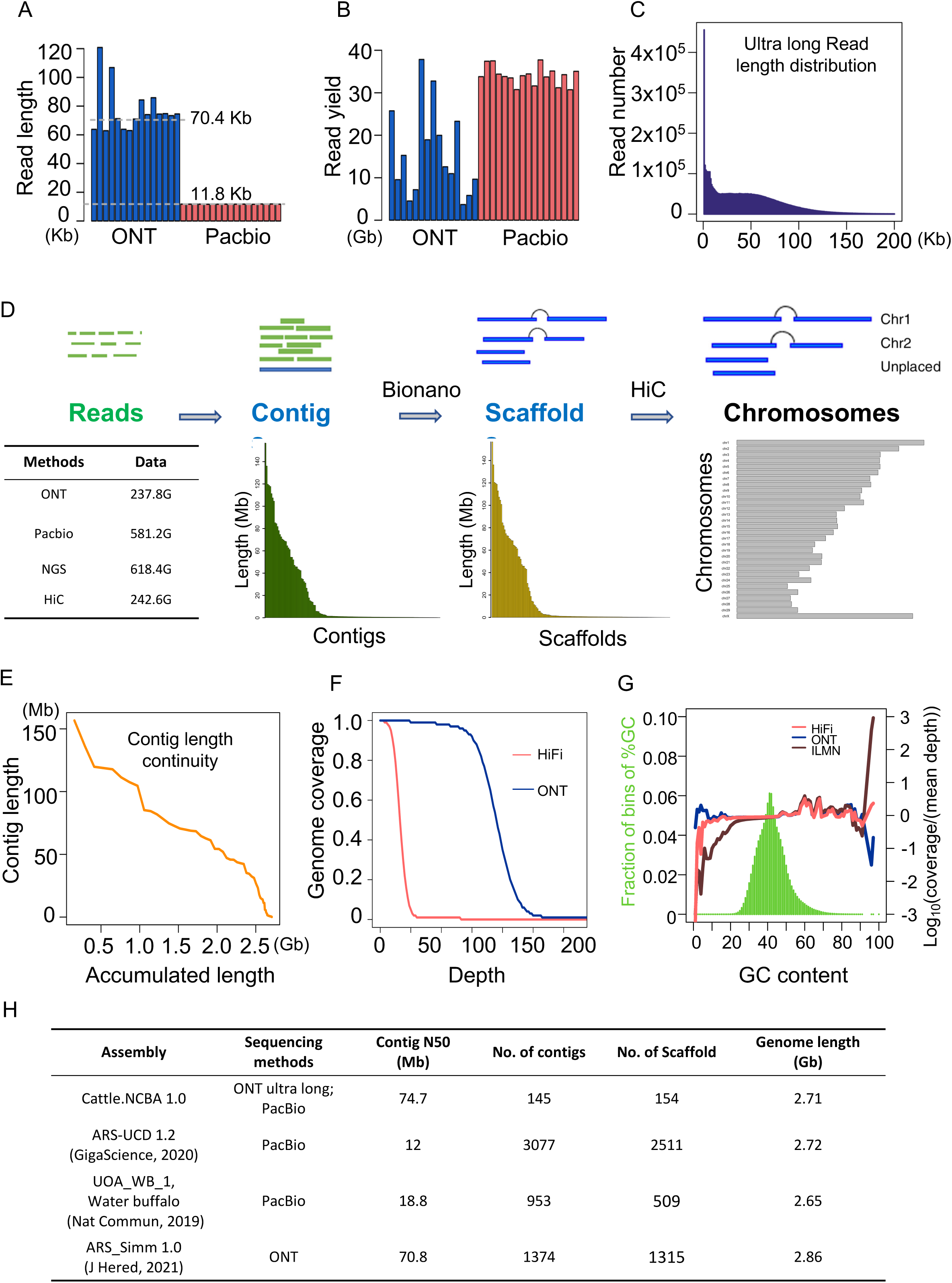
Data Summary of Cattle Genomic Assembly. (A) Read length distributions of ONT ultra-long and PacBio HiFi methods. Each bar indicates a separate sequencing flow cell. (B) Read yield distributions of ONT ultra-long and PacBio HiFi methods. Each bar indicates a separate sequencing flow cell. (C) Length distribution of the total ONT ultra-long reads. (D) Flow chart of the genome assembly pipeline. (E) Continuity of ultra-long reads assembled contigs. (F) Read coverage of NCBA1.0 by ONT ultra-long reads and HiFi reads. (G) Influence of GC content on genome coverage of three sequencing methods. (H) Summary of recently assembled genomes related to bovine.

**Figure. S2.**
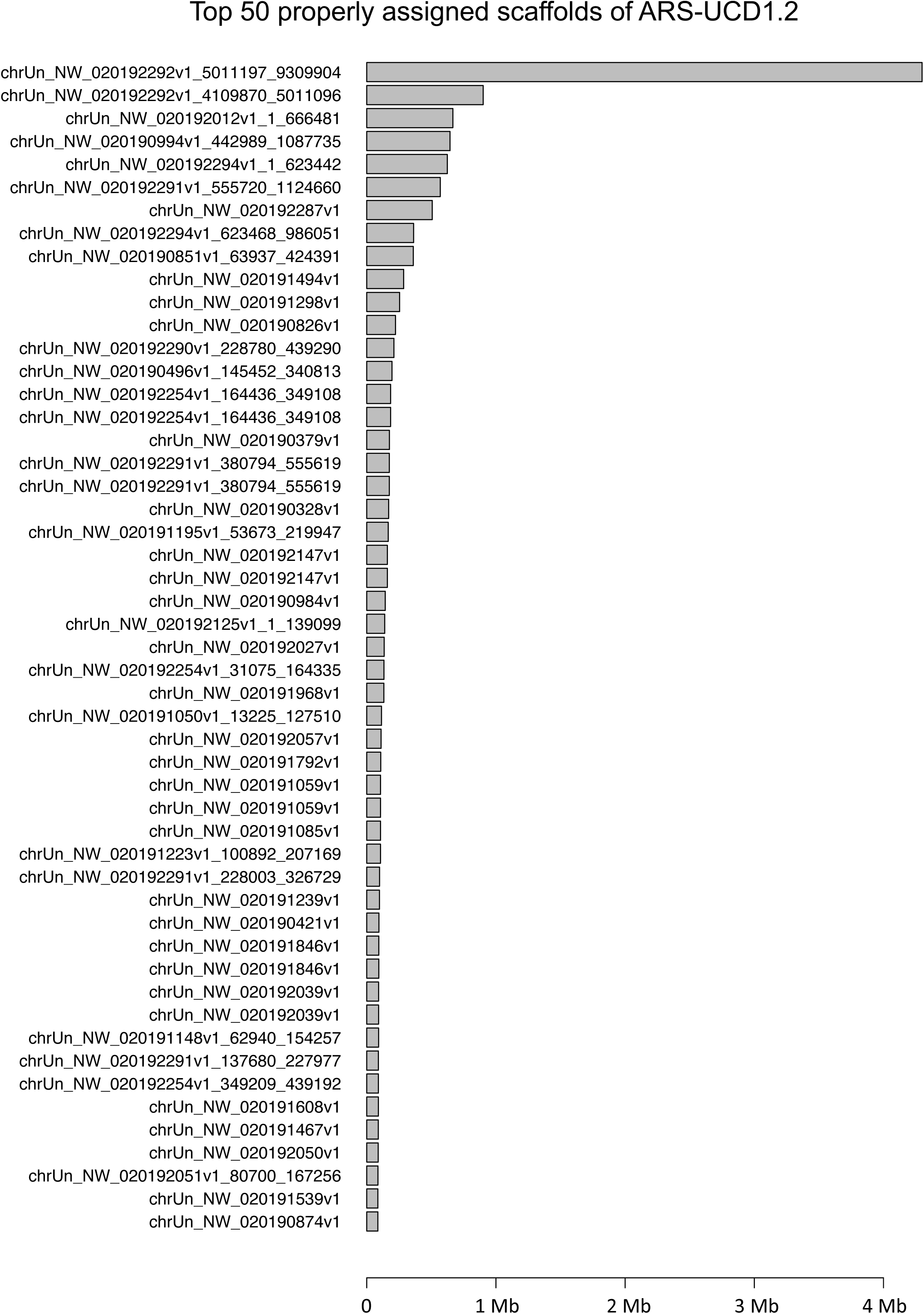
Properly Placed Scaffolds of ARS-UCD1.2 in NCBA1.0. Top fifty scaffolds of ARS-UCD1.2 according to their sequence length were shown. Information of all properly placed scaffolds were stored in Table S7.

**Figure S3.**
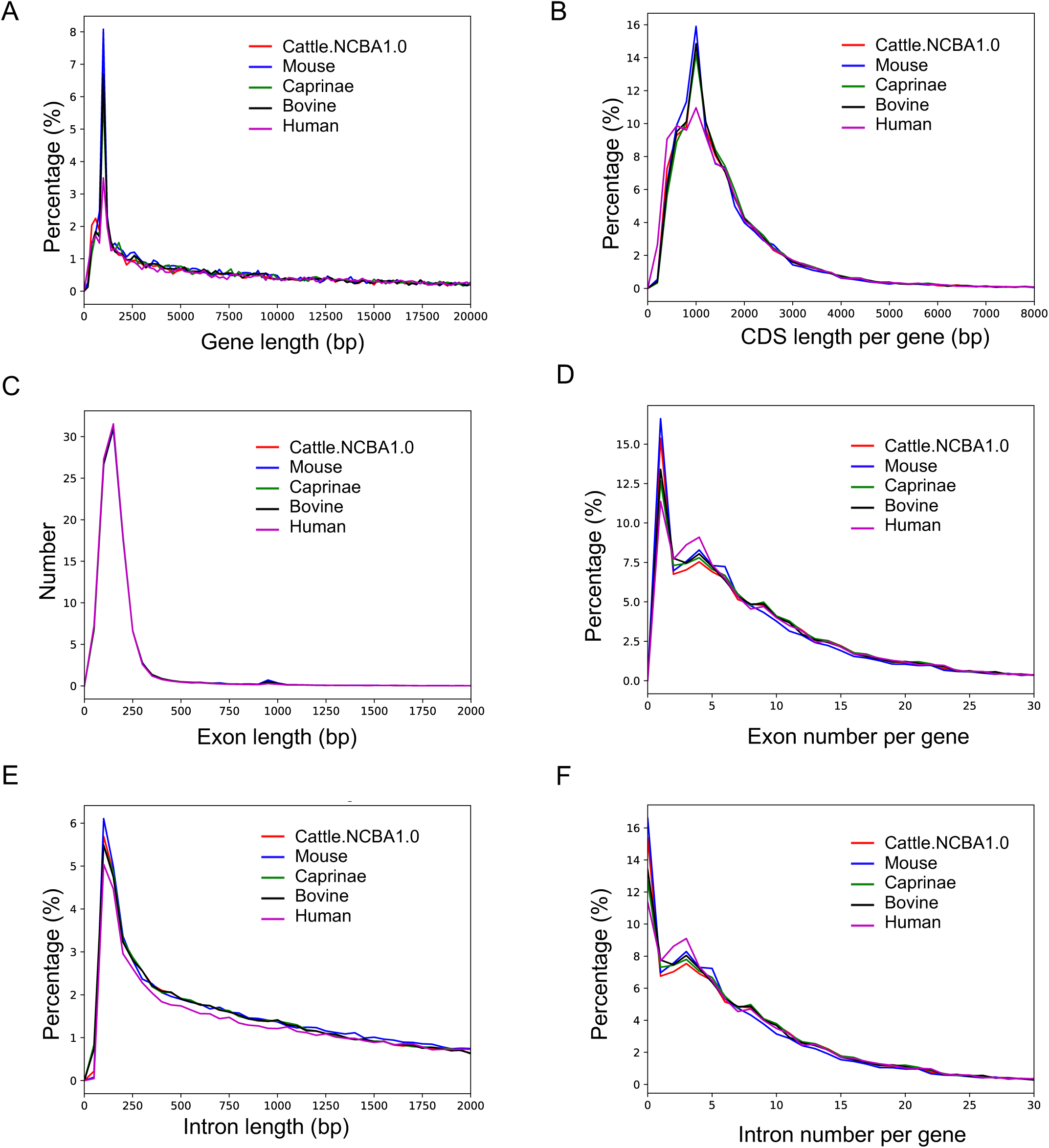
Gene Length Distributions of Five Species. (A) Distribution of the gene lengths of five species. (B) Distribution of the CDS lengths of five species. (C) Distribution of the exon lengths of five species. (D) Distribution of the exon numbers per gene of five species. (E) Distribution of the intron lengths of five species. (F) Distribution of the intron numbers per gene of five species.

**Figure S4.**
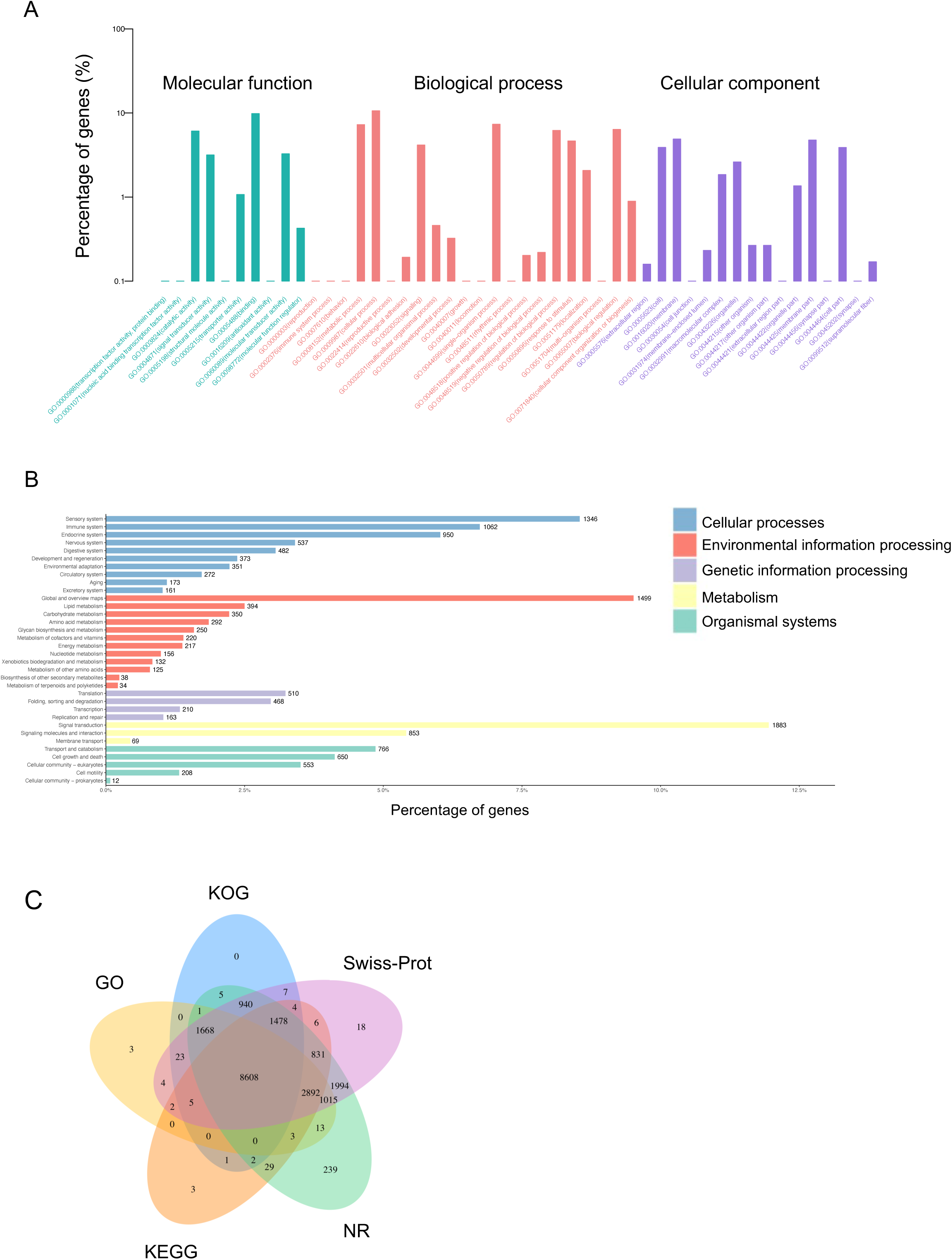
Functional Annotation of Predicted Genes in NCBA1.0. (A) GO annotation of predicted genes. (B) KEGG Ortholog annotation of predicted genes. (C) Venn diagram of genes annotated to KOG, GO, KEGG, NR and Swiss-Prot.

**Figure S5.**
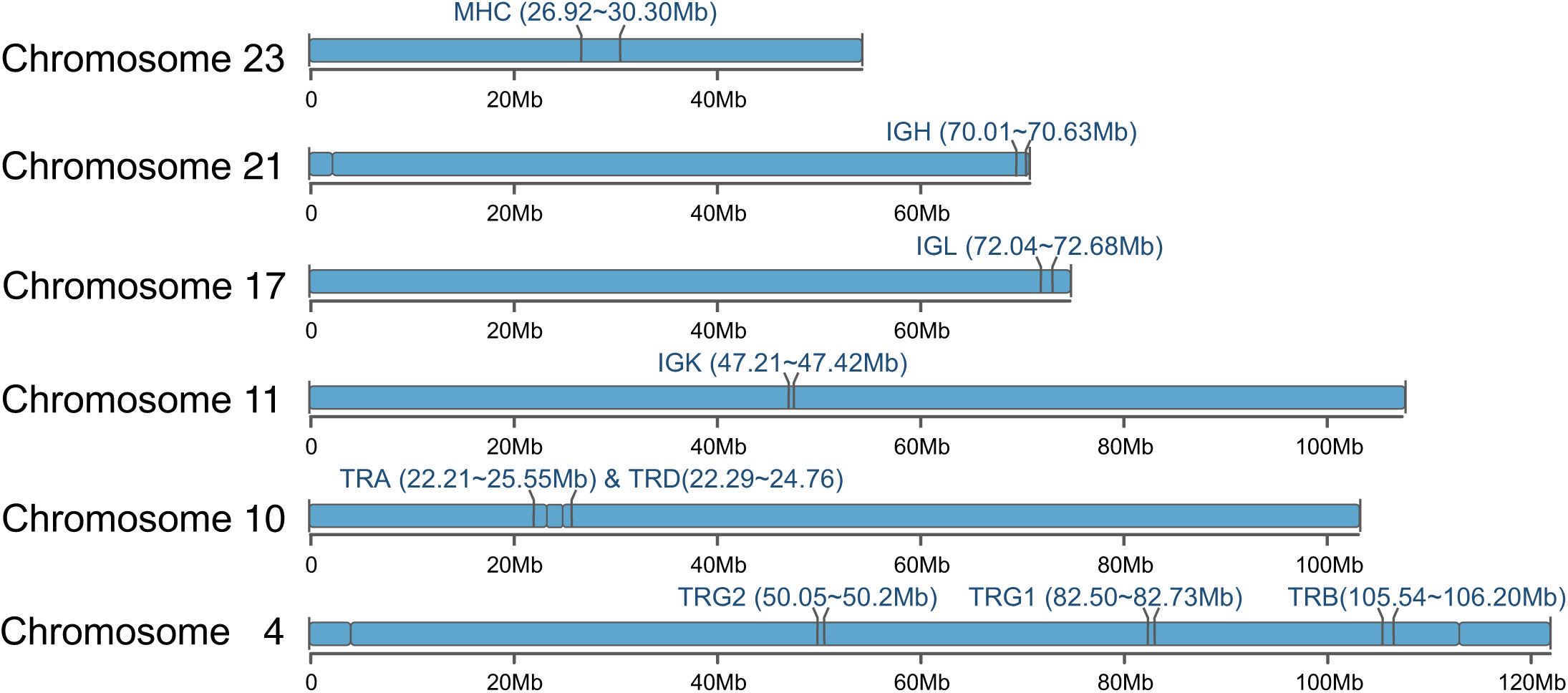
Immune Gene Loci in NCBA1.0 Assembly. The IG, TR and MHC genomic loci dispersed in six chromosomes. The genomic coordinates for each locus were labeled and the gaps between contigs were annotated too. All immune loci were seamlessly assembled except the two gaps within the TRA/TRD region.

**Figure S6.**
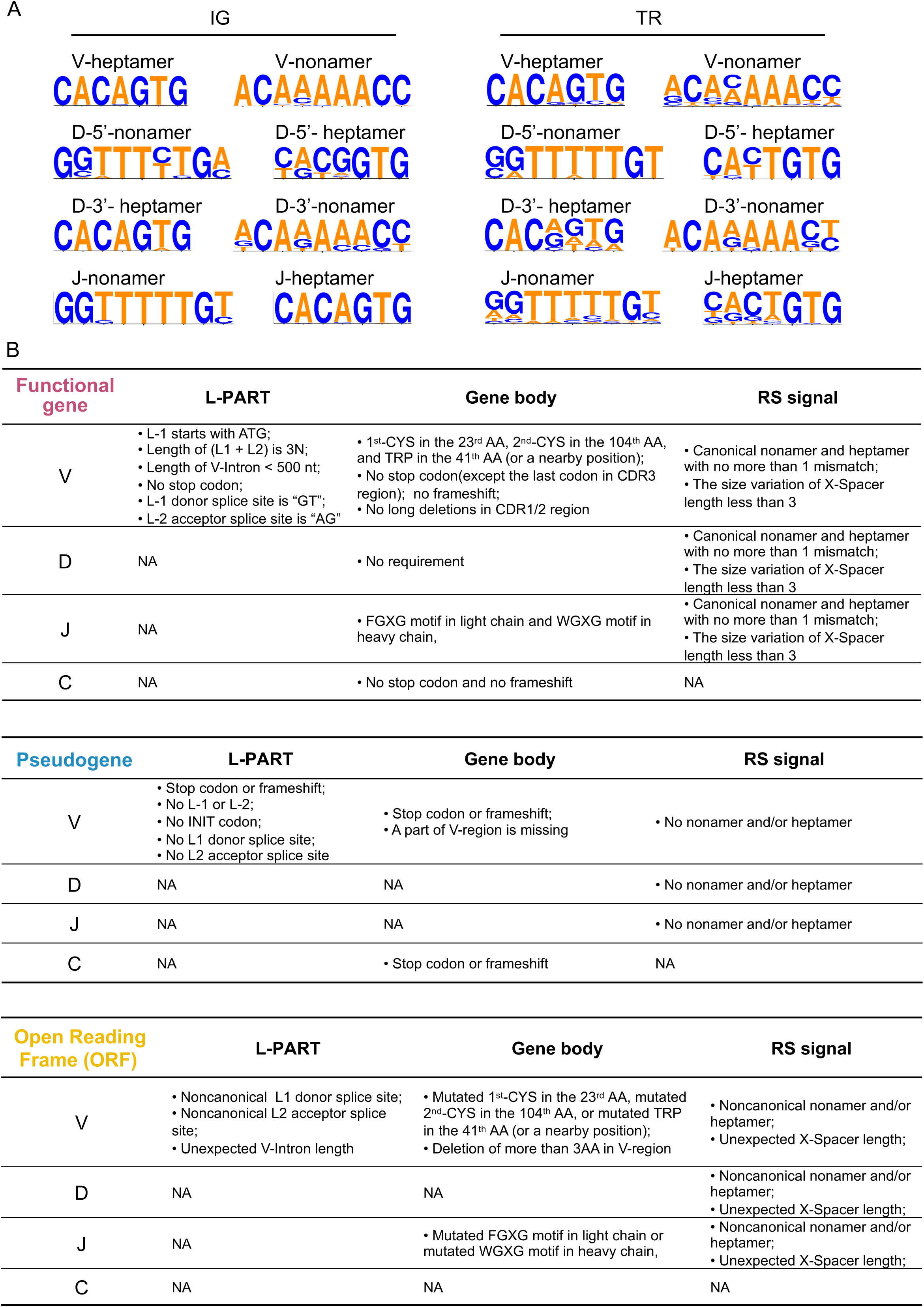
Criteria For Gene Structure and Functionality Annotation, (A) Sequence conservation logos were created with recombination signal sequences of all functional cattle genes from IMGT by WebLogo software. (B) Criteria for determining the functionality of an IG/TR gene. The functionality of an IG/TR gene is defined as functional (F), Open Reading Frame (ORF) or Pseudogene (P) based on the sequence analysis.

**Figure S7.**
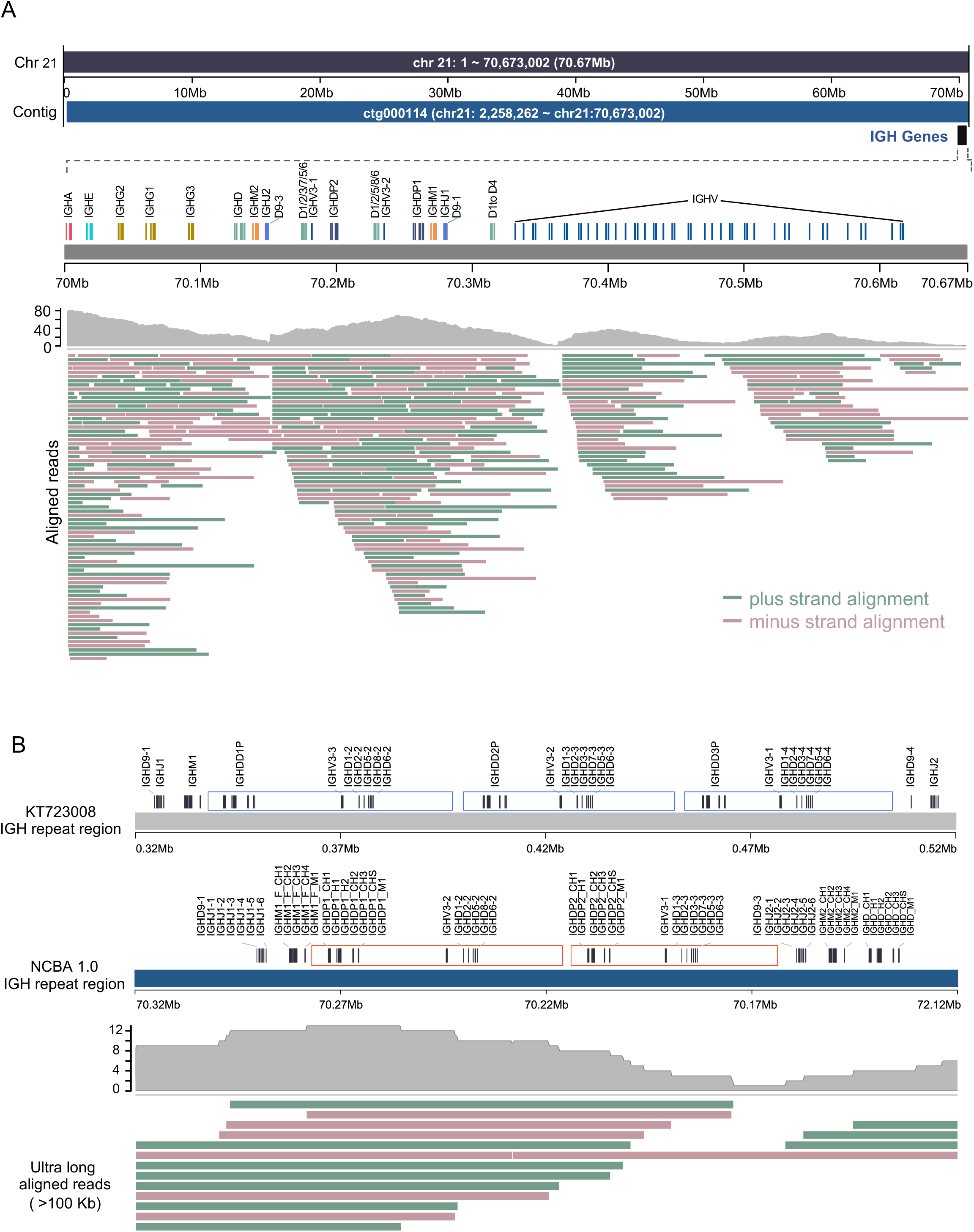
Ultra-long Reads Coverage of IGH Locus. (A) Global view of the IGH locus covered by ONT ultra-long reads. (B) Enlarged tandem repeat regions of IGH locus. The repeated regions were represented as blue and red rectangles in KT723008 and NCBA1.0, respectively.

**Figure S8.**
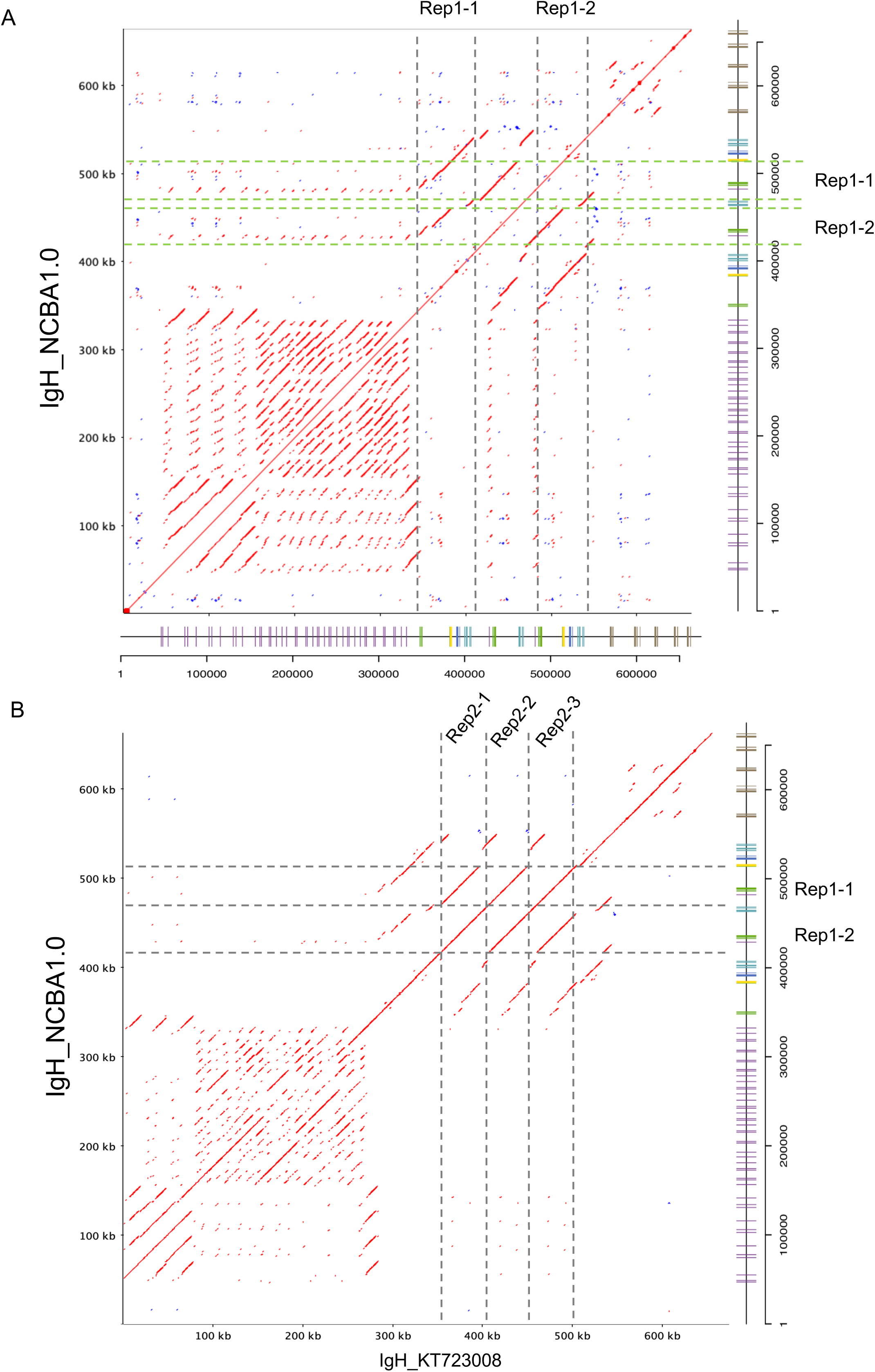
Dot Plots between Two IGH Genomic Sequences. (A) Dot plot of the IGH genomic sequence in NCBA1.0 assembly. The repeated regions were labeled as rep1-1 and rep1-2. (B) Pairwise alignment between IGH in NCBA1.0 and previously reported IGH sequence (KT723008). The three tandem repeats in KT723008 were labeled as rep2- 1, rep2-2 and rep2-3.

**Figure S9.**
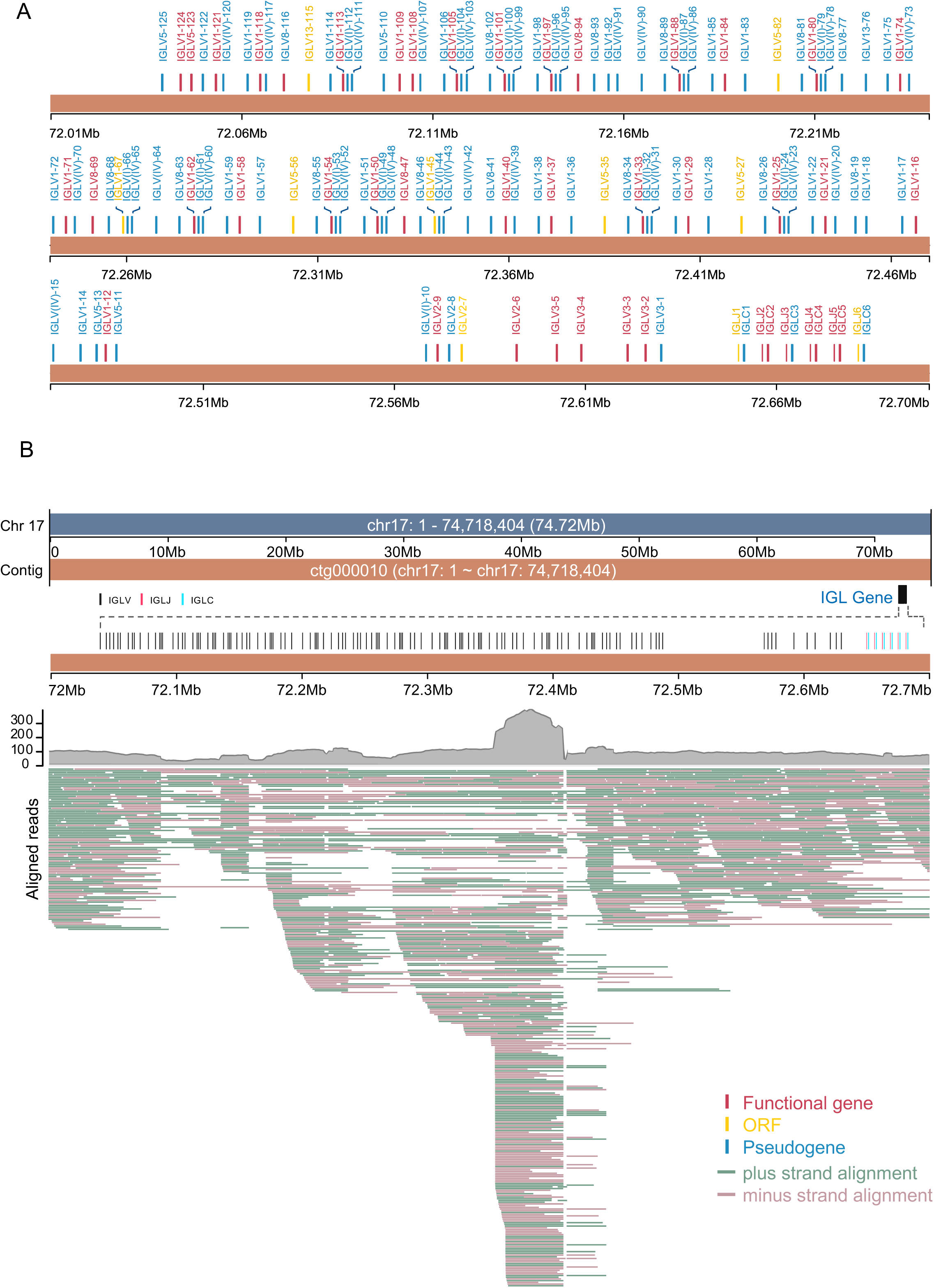
Detailed Annotation Map of IGL Locus. (A) Elaborate gene structures of IGL locus in NCBA1.0 assembly. (B) Read coverage by ONT ultra-long reads in the IGL genomic region.

**Figure S10.**
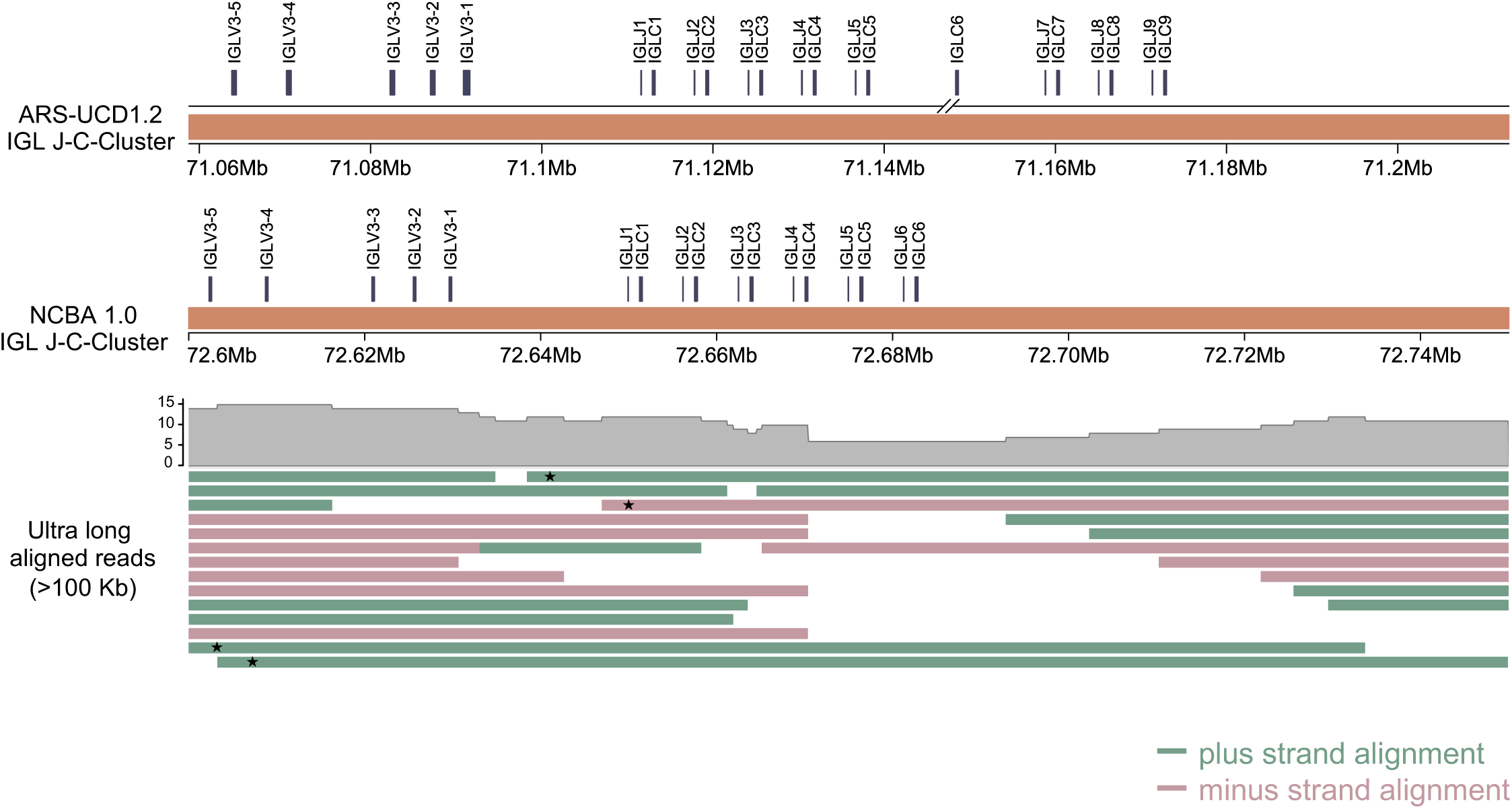
Enlarged Alignment Map of IGL J-C Cluster Region. ONT ultra-long reads that longer than 100 Kb were aligned back to the IGL locus, and four separate ONT ultra-long reads that span over the entire IJL J-C cluster region were labeled with asterisk.

**Figure S11.**
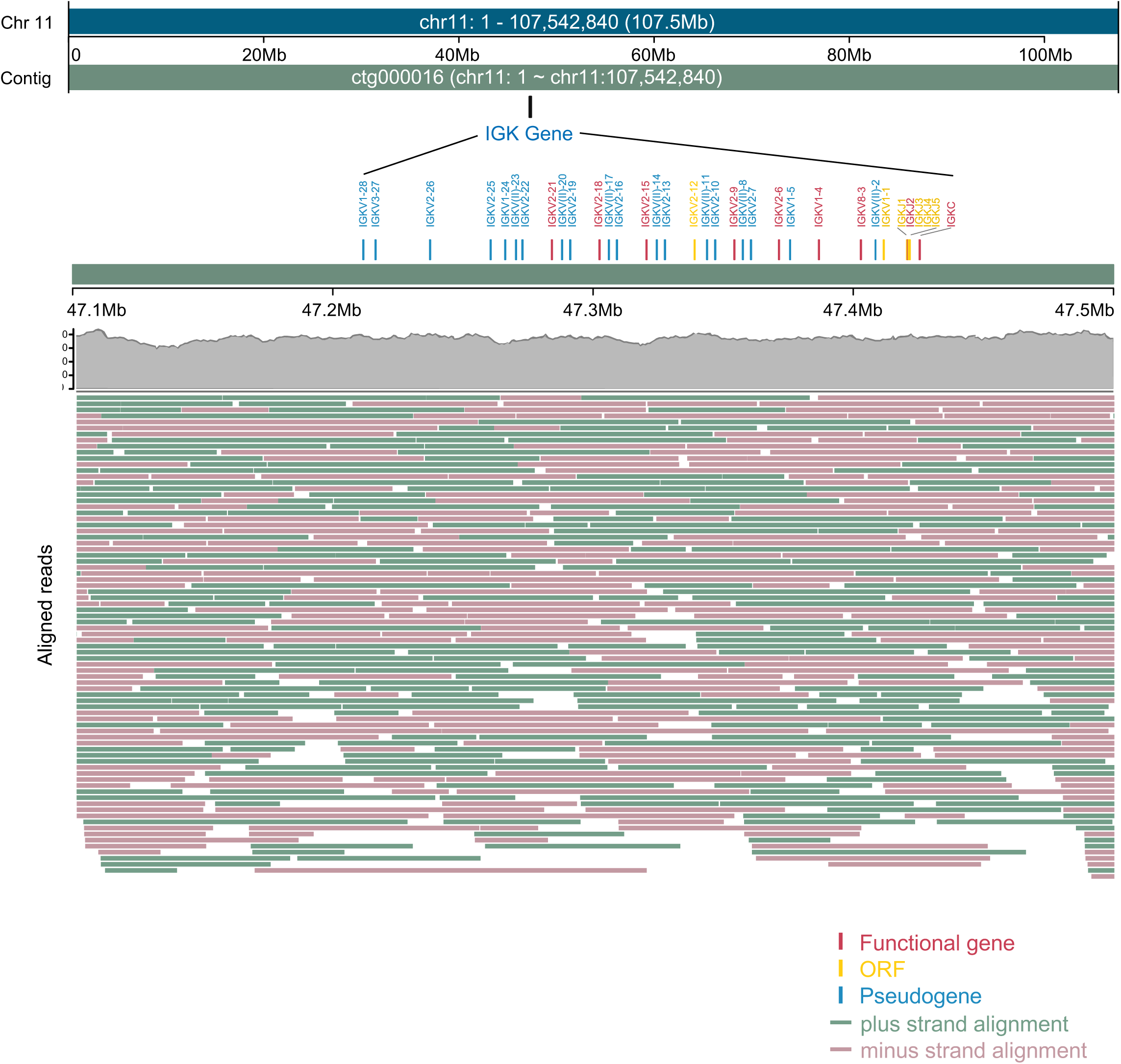
Detailed Annotation Map of IGK Locus. Genomic coordinate and organization of IGK locus were depicted. ONT ultra-long reads that longer than 100 Kb and the mapping to the genomic region were drawn.

**Figure S12.**
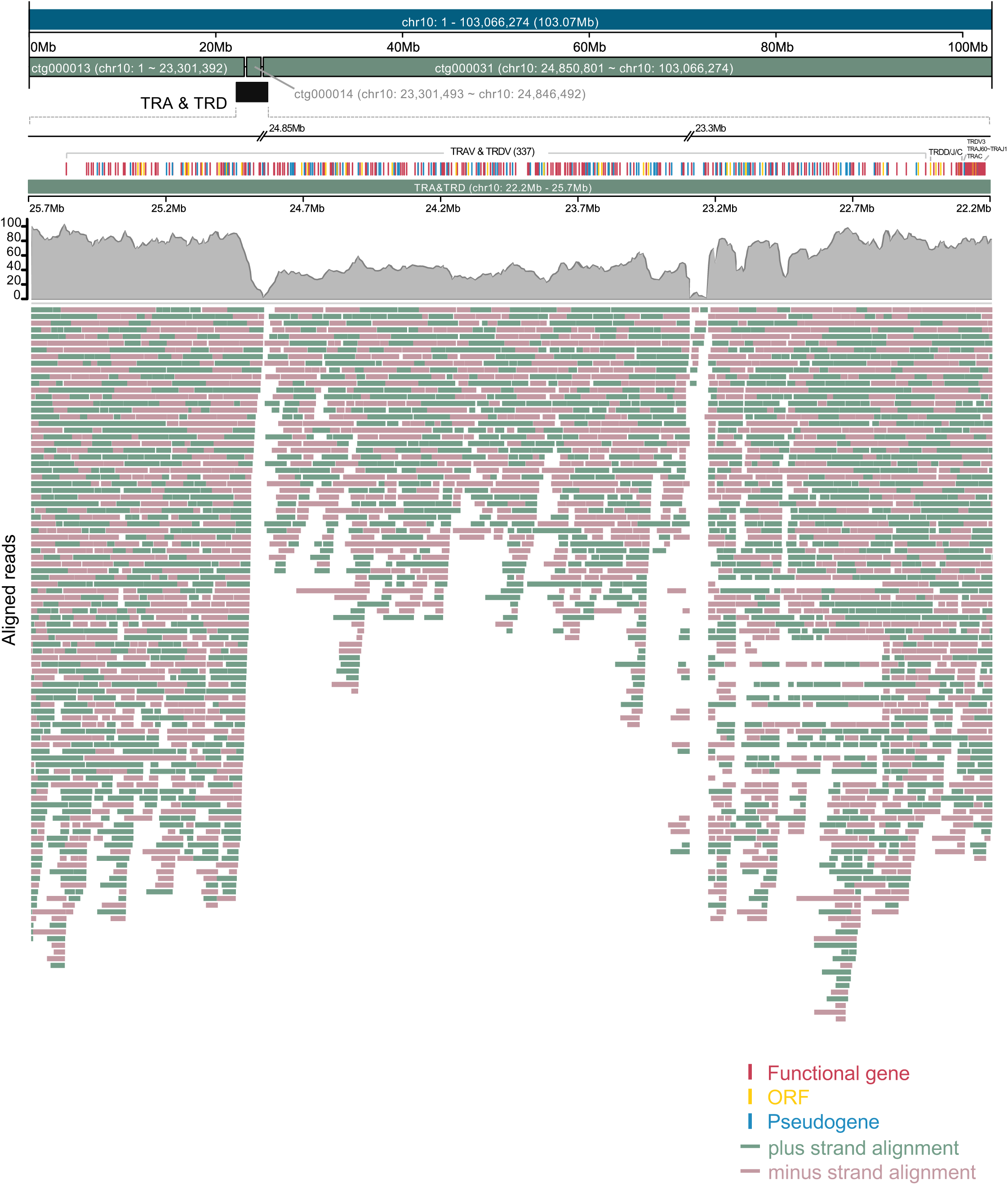
Global Genetic Map of TRA/TRD loci. Genomic coordinate and organization of TRA/TRD loci were annotated. ONT ultra-long reads that longer than 100 Kb and the mapping to the genomic region were drawn. The remaining two gaps within the TRA/D V gene region were depicted too.

**Figure S13.**
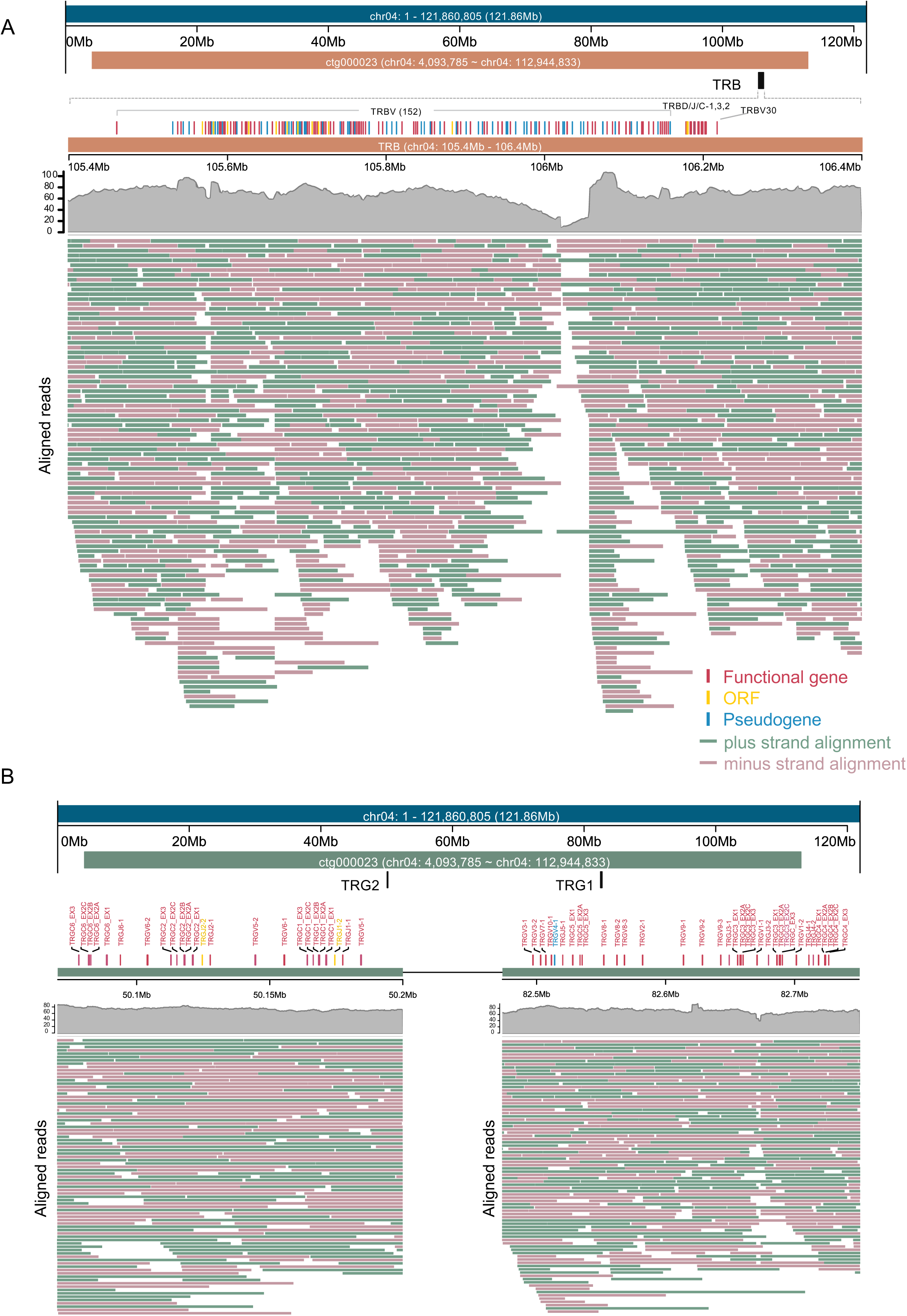
Global Alignment Map of TRB and TRG Loci. (A) Genetic map of TRB locus in chromosome 4. TRB locus resides within contig23 and ONT ultra-long reads that mapped to the genomic region were drawn. (B) Genetic map of TRG locus in chromosome 4. TRG contains two separate gene clusters: TRG1 and TRG2, that are 32 Mb distant away from each other. ONT ultra-long reads that mapped to the genomic region were drawn.

**Figure S14.**
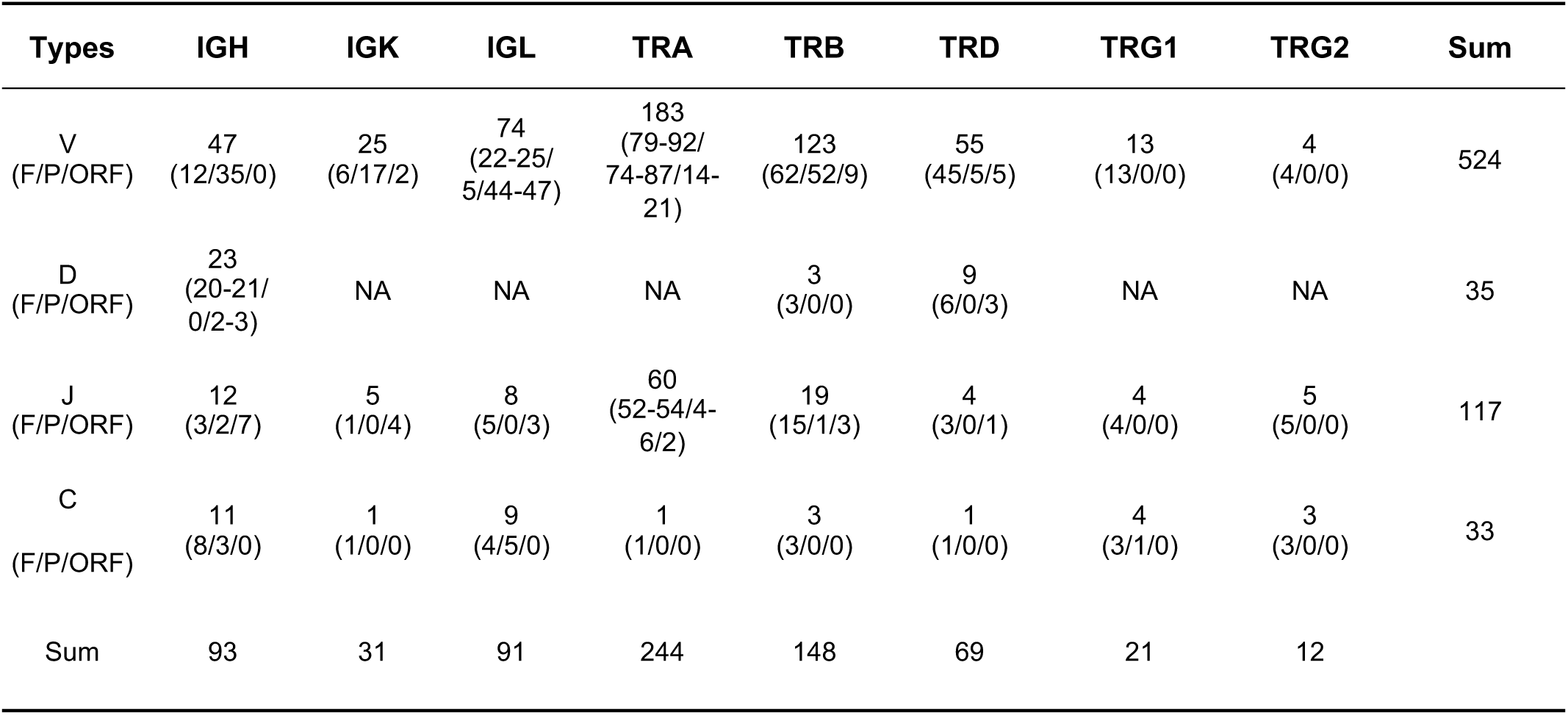
Gene Statistics of Cattle for Each Immune Locus in IMGT. Table was drawn based on cattle gene numbers of IG and TR collected from IMGT database.

**Figure S15.**
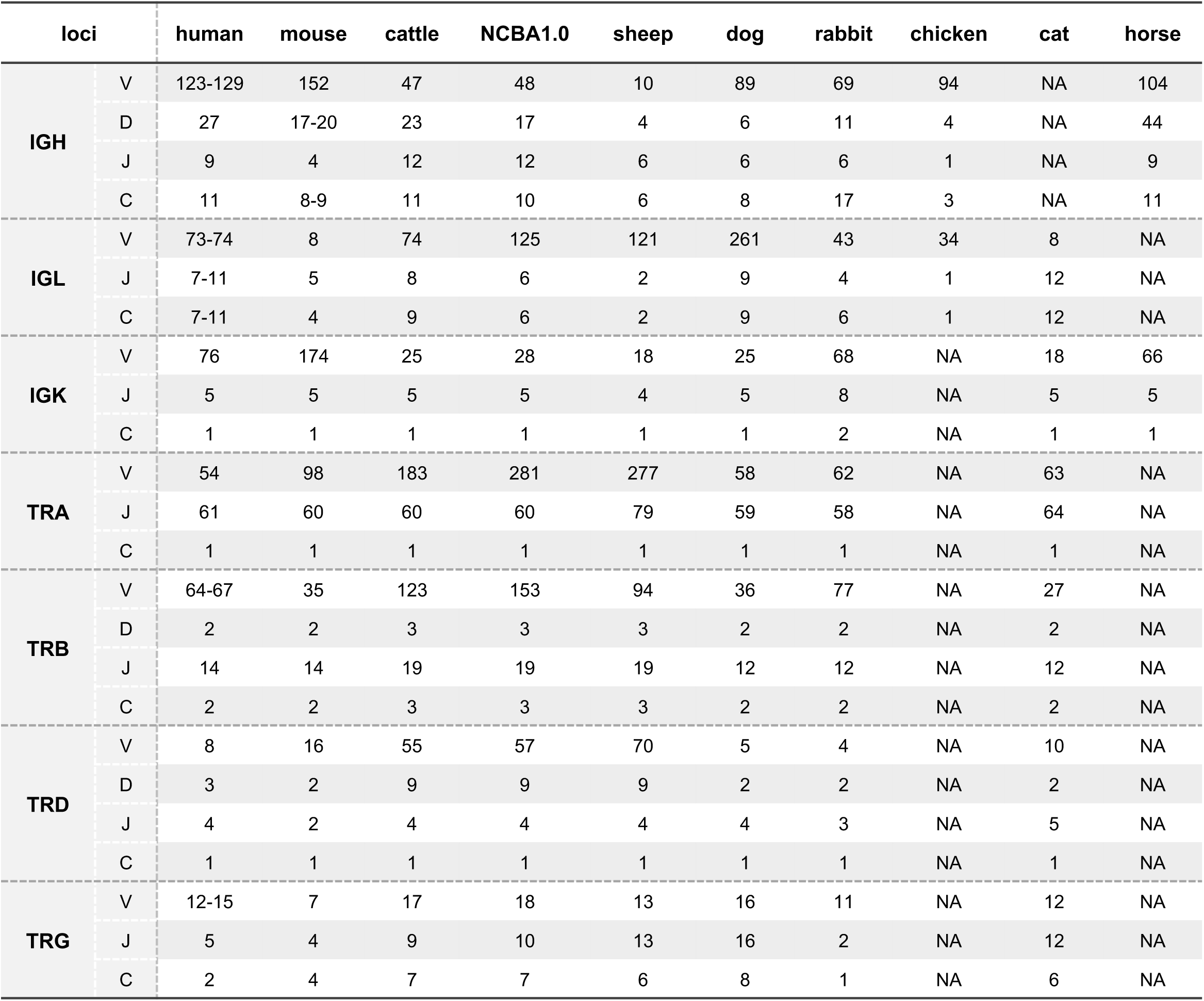
IG and TR Statistics of Different Species in IMGT. Table was drawn based on gene numbers of IG and TR of different species collected from IMGT database.

**Figure S16.**
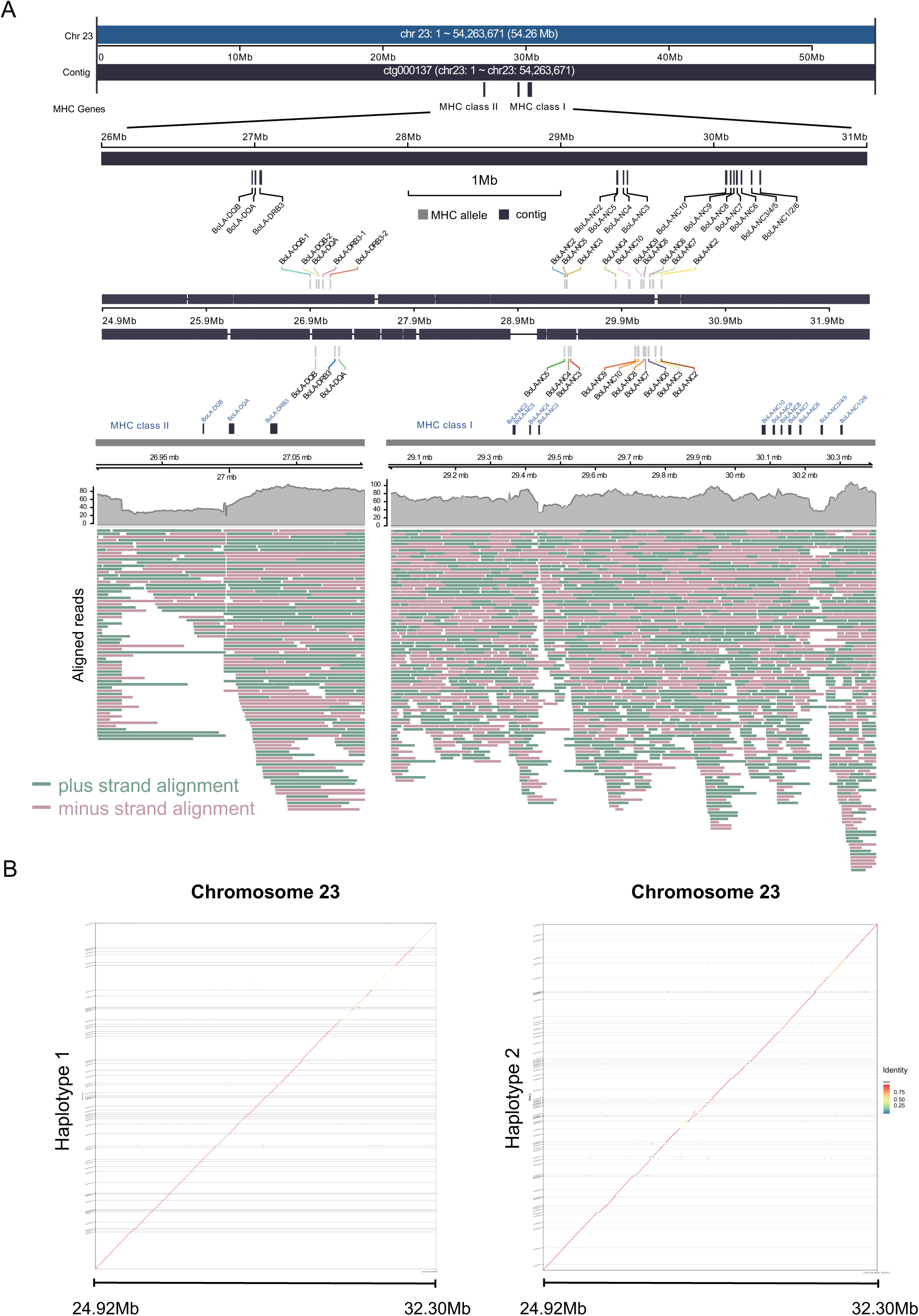
Genomic Assembly and Haplotyping of MHC Locus. (A) Genomic coordinate and annotation of MHC I and MHC II. ONT ultra-long reads that mapped to the genomic region were drawn. (B) Sequence alignments between two haplotigs and the MHC genomic region. Colors indicate the sequence identity of the alignments.

**Figure S17.**
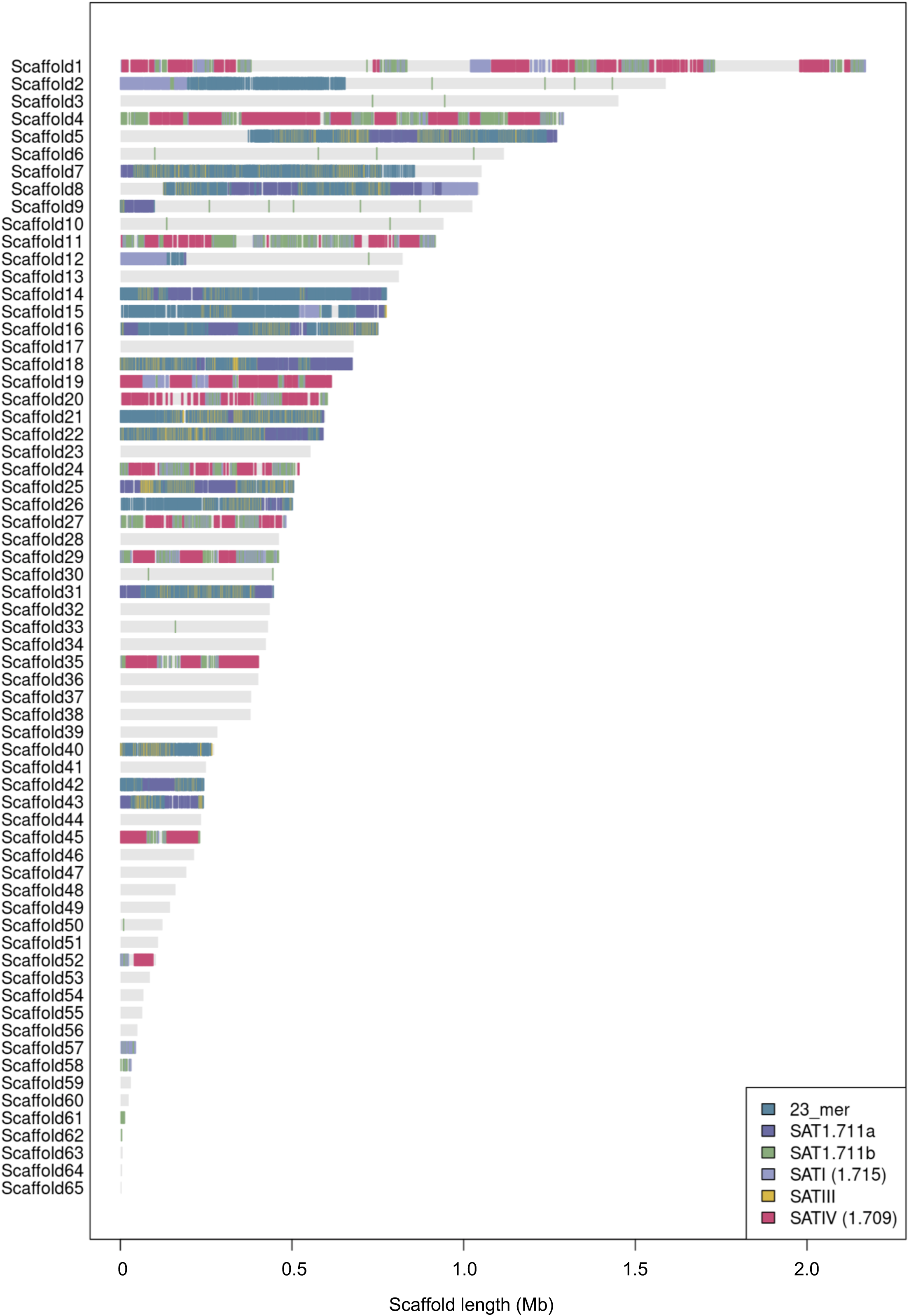
Distributions of satDNAs in Scaffolds. Genomic loci of satDNAs within the scaffolds of NCBA1.0 were drawn.

